# 3′UTR-derived small RNA couples acid resistance to metabolic reprogramming of *Salmonella* within macrophages

**DOI:** 10.1101/2025.06.06.658255

**Authors:** Takeshi Kanda, Fang Liu, Hoda Kooshapour, Sarah Reichardt, Maolin Wang, Philippe Icyishaka, Nozomu Obana, Alexander J. Westermann, Yanjie Chao, Masatoshi Miyakoshi

## Abstract

Acid resistance is crucial for enterobacteria to withstand host acidic environments during infection, including the gastrointestinal tract and macrophage phagosomes. A key acid resistance mechanism of the facultative intracellular pathogen *Salmonella* is expression of the arginine decarboxylase AdiA. While AdiA confers acid resistance via an H^+^-consuming reaction, we discover that the 3′-untranslated region (UTR) of *adiA* mRNA is processed by RNase E into a regulatory small RNA, AdiZ. Through RNA-RNA interactome profiling and transcriptomic analysis, followed by *in vitro* structural probing and *in vivo* validations, we demonstrate that AdiZ directly base-pairs with and negatively regulates *ptsG*, *pykF,* and *dmsA* mRNAs involved in glucose uptake, glycolysis, and anaerobic respiration, respectively. Intriguingly, AdiZ is induced and facilitates *Salmonella* survival within macrophages, where acidic and hypoxic stresses prevail. Thus, simultaneous expression of AdiA and AdiZ from a single mRNA ties arginine-dependent acid resistance to metabolic reprogramming of *Salmonella* in the host intracellular niches.

**HIGHLIGHTS:**

- *Salmonella adiA* transcript expresses both acid resistance enzyme and sRNA AdiZ
- AdiZ derived from the *adiA* 3′UTR is induced under an acidic and anaerobic condition
- AdiZ base-pairs with mRNAs to modulate glucose metabolism and anaerobic respiration
- AdiZ promotes *Salmonella* survival in acidic and hypoxic macrophage vacuoles

**GRAPHICAL ABSTRACT:** 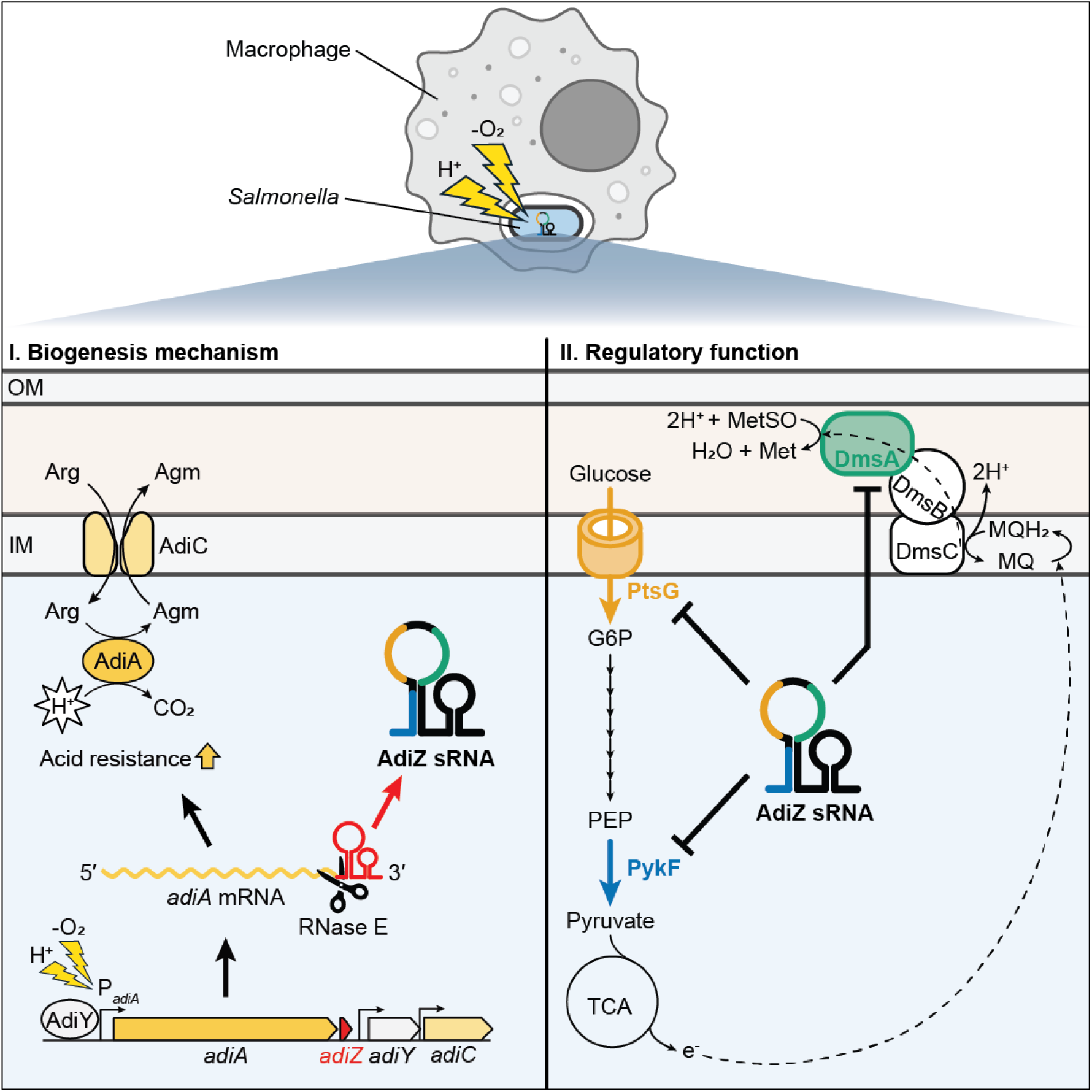

## INTRODUCTION

*Salmonella enterica* (hereafter referred to as *Salmonella*) is a globally important pathogen responsible for over 100 million infections annually (1). This bacterium is primarily transmitted through the fecal-oral route, with a higher prevalence in regions without proper sanitation or access to clean drinking water (2, 3). During infection, *Salmonella* experiences different acidic environments; extracellularly, throughout the gastrointestinal tract (4), and intracellularly, during its proliferation inside the mildly acidic *Salmonella*-containing vacuole (SCV) within host macrophages (5). For *Salmonella* to adapt to acidic environments, key enzymes to its survival in acidic environments are inducible amino acid decarboxylases. The *Salmonella* genome encodes three acid-inducible decarboxylases, AdiA, CadA, and SpeF, which catalyze the H⁺-consuming conversion from arginine, lysine, and ornithine to agmatine, cadaverine, and putrescine, respectively. The cognate antiporters, AdiC, CadB, and PotE, transport the polyamines out of the cell in exchange for their respective amino acids (Figure 1A) (6, 7). The CadA/CadB and SpeF/PotE systems are required to counteract growth inhibition by short-chain fatty acids (SCFAs) produced by the gut microbiota in the large intestine (8). Among these acid resistance systems, the arginine-dependent system is the most effective in protecting *Salmonella* under anoxic conditions against an extreme acidic stress (pH 2.3) (9). The transcription of *adiA* is activated under acidic and anaerobic conditions by an AraC/XylS family transcriptional regulator, AdiY (10), which is encoded between *adiA* and *adiC* genes on the same strand (Figure 1A). The *adiA* gene contains an intrinsic terminator, which attenuates the transcription of downstream genes (11) and whose transcript binds the RNA chaperone Hfq (12). The 3′-untranslated region (UTR) of the monocistronic *adiA* mRNA accumulates as a 95-nt small RNA (sRNA), annotated as STnc1180 (13–15). However, the mechanism of STnc1180 biogenesis and the potential regulatory function of this sRNA were unclear.

**Figure 1.**
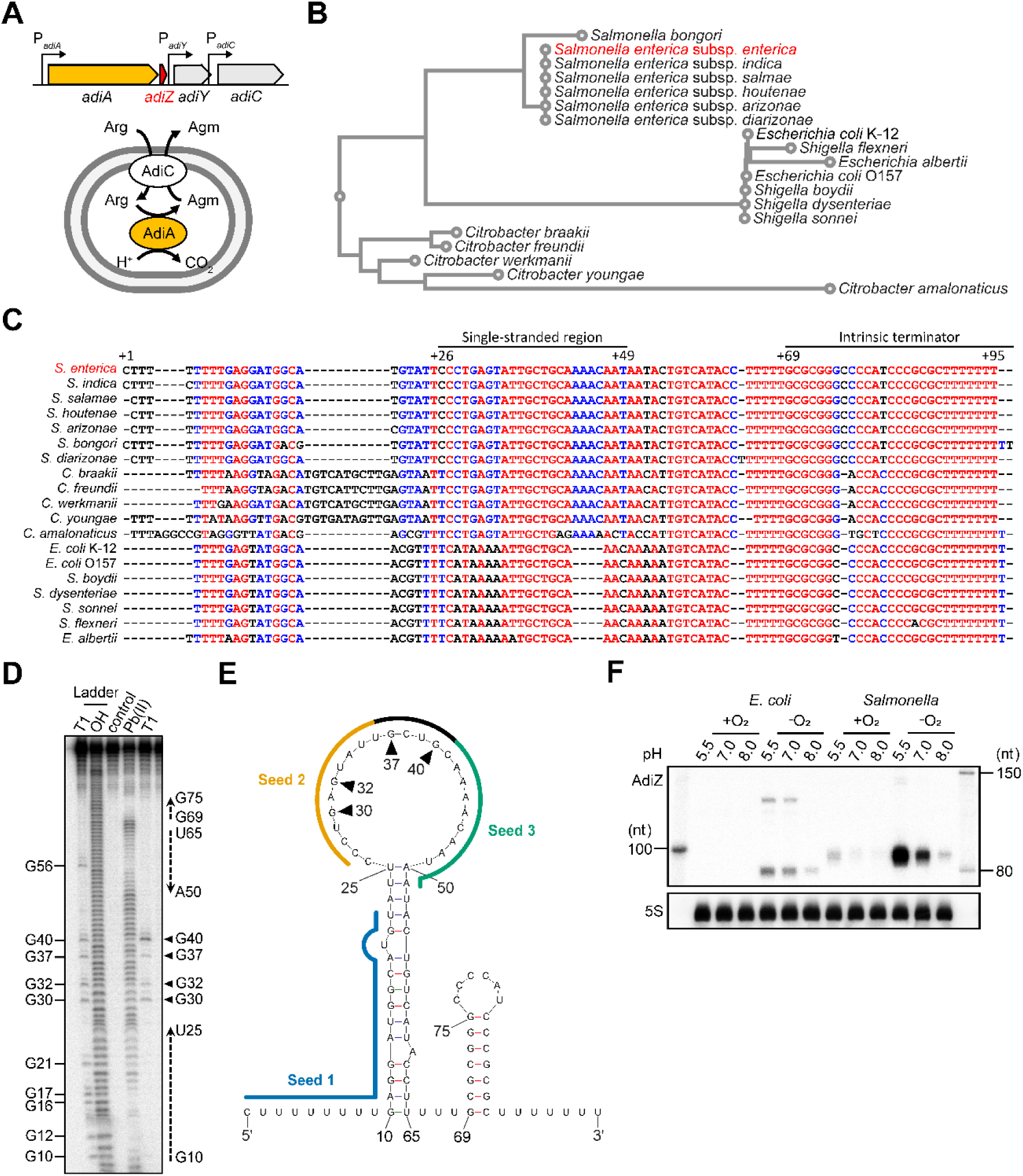
Conservation, secondary structure, and expression profile of AdiZ sRNA. (A) Schematic representation of the *adiAYC* cluster (top) and the AdiA/C acid resistance system (bottom). AdiA: arginine decarboxylase; AdiY: transcriptional activator of *adiA* and *adiC*; AdiC: arginine:agmatine antiporter. AdiA consumes intracellular H⁺ through arginine decarboxylation. *adiZ* is located 12 bp downstream of the *adiA* stop codon in *Salmonella*. (B) Phylogenetic tree of *adiZ* constructed using MAFFT (89). (C) Multiple sequence alignment of *adiZ* performed using MultAlin (90). Fully, partially and poorly conserved nucleotides are indicated in red, blue, and black, respectively. The single-stranded region in *Salmonella* AdiZ and the conserved intrinsic terminator are indicated. (D) *In vitro* structure probing of 5′end-labeled *Salmonella* AdiZ. “T1” and “OH” ladders represent digestion under denaturing conditions with RNase T1 (lane 1, cleaving unpaired G residues) and alkali treatment (lane 2, cleaving at all positions), respectively. Lane 3 (“control”) shows untreated AdiZ, while lanes 4 and 5 depict cleavages induced by lead(II) acetate (cleaving single-stranded regions) and RNase T1 under native conditions, respectively. (E) Secondary structure of *Salmonella* AdiZ predicted from *in vitro* structure probing data and computational analysis using mfold (37). Unpaired G residues identified in (D) are marked with arrowheads. Three seed regions as defined in Figure 4A are indicated. The region shared by seed 2 and seed 3 is shown in black. (F) AdiZ expression is highest under acidic (pH 5.5) and anaerobic conditions in both *E*. *coli* and *Salmonella*. Bacteria were cultured at pH 7.0 under aerobic or anaerobic conditions until the late exponential phase. The culture pH was shifted to acidic (pH 5.5), alkaline (pH 8.0), or maintained at neutral pH (pH 7.0). Total RNA was extracted 30 min after the pH shift and analyzed by northern blot. AdiZ was detected using probes MMO-1468 and MMO-1469 for *E*. *coli* and *Salmonella*, respectively. 5S rRNA detected by MMO-1056 served as a loading control.

Hfq-dependent sRNAs typically repress translation of target mRNAs by base-pairing near the ribosome-binding site, thereby blocking ribosome access, which often accompanies the degradation of the target mRNA (16–18). sRNAs derived from mRNA 3′UTRs similarly function as canonical Hfq-dependent sRNAs (19–22). Given that the nucleotide sequences in the 3′UTRs are highly conserved among bacterial species, their complementarity to the other mRNAs often in the 5′UTRs implicates profound links in various biological processes such as central metabolism (23–27), anaerobic respiration (28), and membrane integrity (29). Recent advances in RNA-seq methodologies have expanded our knowledge of bacterial RNA-RNA interactions (30–32), but most of them await further investigation.

In this study, we extend the physiological roles of the *adi* locus beyond its well-established function in acid resistance. We demonstrate that STnc1180, which we rename to AdiZ, is released by RNase E from the 3′UTR of *adiA* mRNA, whose expression is induced under an acidic and anaerobic condition. By analyzing both transcriptome and RNA-RNA interactome of *Salmonella* grown under specific conditions, we identify *pykF*, *ptsG*, and *dmsA* as direct targets of AdiZ. These genes are critical for glycolysis (33), glucose uptake (34), and anaerobic respiration (35), respectively, thereby linking acid resistance with key metabolic processes of *Salmonella* within macrophages. AdiZ specifically interacts with these mRNAs using three distinct seed sequences. These post-transcriptional regulations support *Salmonella* proliferation within the acidic and anoxic environment inside macrophages.

## RESULTS

### AdiZ is expressed under an acidic and anaerobic condition

To investigate the genetic distribution of AdiZ, we compared the sequences of the *adiA* 3′UTR across members of the *Enterobacteriaceae* family, where *adiA* is predominantly conserved (36). The *adiZ* homologs were grouped into three distinct clusters, *Escherichia*-*Shigella*, *Citrobacter*, and *Salmonella* (Figure 1B). In all clusters, the Hfq-binding module, composed of the intrinsic terminator preceded by a U-rich sequence (12), was nearly perfectly conserved (Figure 1C). In contrast, the other sequences of AdiZ are relatively variable and exhibit cluster-specific characteristics. *In vitro* structure probing using lead(II) and RNase T1 combined with *in silico* prediction by mfold (37) revealed that *Salmonella* AdiZ contains a stem-loop with a 24-nt single-stranded region, which would be available for base pairing with target mRNAs (Figures 1D and 1E).

To profile the expression of AdiZ, we performed northern blot analysis. *Salmonella* and *E. coli* were cultured in neutral LB (pH 7.0) under either aerobic or anaerobic conditions until the late exponential phase. The culture was then acidified (pH 5.5), alkalinized (pH 8.0), or kept neutral (pH 7.0). Total RNA was extracted 30 min after the pH shift and analyzed by northern blot. In *Salmonella*, a single band of ∼90-nt transcript was detected, whereas in *E. coli*, two distinct bands corresponding to ∼80 and ∼130 nt were observed (Figure 1F). The signal intensities of AdiZ isoforms were highest at pH 5.5 under anaerobic conditions, and the expression pattern closely mirrored that of the parental *adiA* mRNA, which was quantified in parallel by qRT-PCR (Figure S1A).

### AdiZ is derived from the 3′UTR of *adiA* mRNA via processing by RNase E

3′UTR-derived sRNAs are either expressed as primary transcripts driven from internal promoters or endonucleolytically processed from their parental mRNAs (19). To distinguish between these biogenesis pathways for AdiZ, we first assessed the role of AdiY, a transcriptional activator required for *adiA* expression (10). Total RNA was extracted from a *Salmonella* Δ*adiY* mutant and the wild-type strain anaerobically grown at pH 5.5. The levels of AdiZ and its parental *adiA* mRNA were analyzed by northern blot and qRT-PCR, respectively. In the Δ*adiY* mutant, the *adiA* mRNA levels were reduced to ∼1/20 of those in the wild type (Figure 2A). Correspondingly, AdiZ expression was abolished in the Δ*adiY* mutant, indicating that AdiY is indispensable for the transcription of both *adiA* and *adiZ*. Notably, expression of *adiY* was induced under anaerobic conditions but not by acidic treatment (Figure S1B). These results suggest that, under anaerobic and acidic conditions, oxygen depletion triggers AdiY expression, then driving transcription of *adiA* and *adiZ*, which is further enhanced in response to acidic signals.

**Figure 2.**
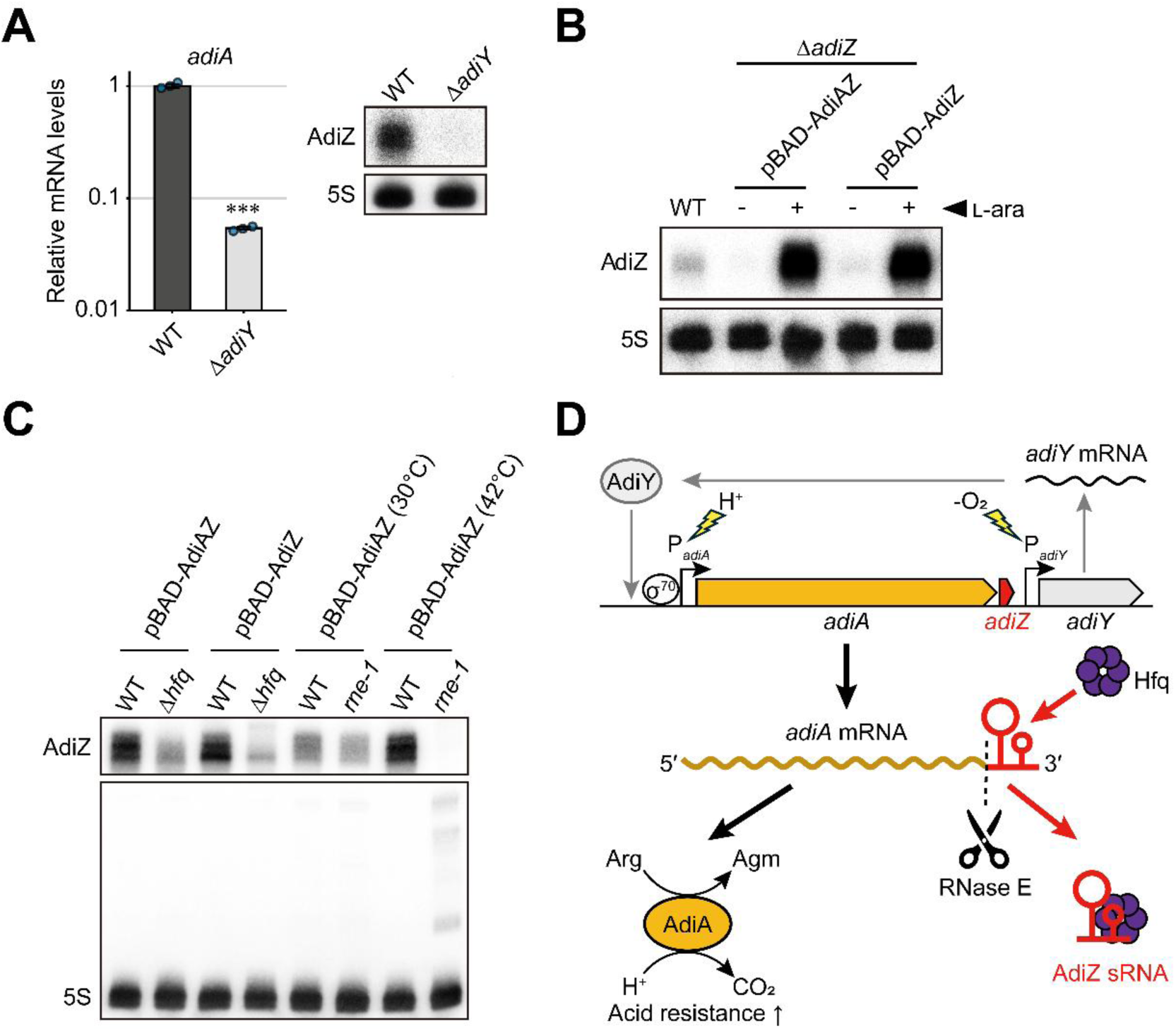
Biogenesis of *Salmonella* AdiZ from the 3′UTR of *adiA* mRNA. (A) Transcriptional activator AdiY is essential for the expression of *adiA* and *adiZ*. Wild-type *Salmonella* and the Δ*adiY* strain were cultured in LB at pH 5.5 under anaerobic conditions. Total RNA was analyzed by qRT-PCR to quantify *adiA* mRNA and by northern blot to detect AdiZ sRNA. qRT-PCR results were normalized to *rpoB* mRNA levels and are presented as mean ± standard error from three independent experiments (n = 3). Statistical significance was assessed using a two-tailed Student’s t-test (***p < 0.001). (B) AdiZ transcription depends on the promoter of *adiA*. The Δ*adiZ* strain harboring either pBAD-AdiAZ or pBAD-AdiZ was cultured in LB at pH 5.5 under anaerobic conditions. At the late exponential phase, 0.2% L-arabinose was added to half of the cultures. After 10 min, total RNA was extracted and analyzed by northern blot. As an endogenous control, total RNA from wild-type *Salmonella* cultured under the same conditions without L-arabinose was included. (C) Hfq and RNase E are required for AdiZ expression. Wild-type and Δ*hfq* strains both with the Δ*adiZ* background harboring either pBAD-AdiAZ or pBAD-AdiZ were cultured in LB at 37°C. At the late exponential phase, 0.2% L-arabinose was added, and the cells were incubated for 10 min. For wild-type and *rne-1* (temperature-sensitive RNase E mutant) strains both in the Δ*adiZ* background harboring pBAD-AdiAZ, cells were cultured at 30°C until the late exponential phase, then either shifted to 42°C or kept at 30°C for 30 min. 0.2% L-arabinose was added, and cells were further incubated for 10 min. Total RNA was analyzed by northern blot. As anticipated, precursors of 5S rRNA accumulated in the *rne-1* mutant at 42°C. (D) Model for AdiZ biogenesis. AdiZ was detected using the probe MMO-1469 and 5S rRNA detected by MMO-1056 served as a loading control in (A), (B), and (C).

Next, we constructed two plasmids: pBAD-AdiAZ, encompassing the *adiA* transcription start site to the *adiZ* terminator, and pBAD-AdiZ, spanning the *adiA* stop codon to the same downstream region, both under the control of the arabinose-inducible P_BAD_ promoter. A *Salmonella* Δ*adiZ* mutant harboring either plasmid was anaerobically grown at pH 5.5. In the presence of arabinose, AdiZ was strongly upregulated in both constructs compared to the endogenous level (Figure 2B). However, in the absence of arabinose, AdiZ expression was negligible, indicating the absence of an internal promoter and supporting the hypothesis that AdiZ is derived from the parental *adiA* mRNA.

The RNA chaperone Hfq is critical for the stability and functionality of numerous sRNAs in *Salmonella* (16), and endoribonuclease RNase E plays a central role in sRNA processing and maturation (38). AdiZ has been previously identified in Hfq co-immunoprecipitation (coIP) assays (14, 39), and TIER-seq (transient inactivation of RNase E followed by sequencing) revealed RNase E cleavage sites near its 5′end (38). To investigate the involvement of these proteins in AdiZ expression, we pulse-expressed either the *adiA-adiZ* region or *adiZ* alone in the Δ*hfq* mutant and the temperature-sensitive RNase E mutant (*rne-1*), respectively. In the Δ*hfq* mutant, AdiZ levels from both constructs were significantly reduced compared to the wild type (Figure 2C). In the *rne-1* mutant, AdiZ expression was comparable to wild-type levels at a permissive temperature (30°C) but was completely abolished by heat inactivation of RNase E at 42°C. These results show that AdiZ is cleaved from the *adiA* transcript by RNase E and is stabilized through its interaction with Hfq (Figure 2D).

### RNA-seq-based approaches reveal putative target mRNAs of AdiZ

To uncover potential target RNAs post-transcriptionally regulated by AdiZ, we first performed RNA-seq analysis following AdiZ pulse expression. A *Salmonella* Δ*adiZ* mutant harboring pBAD-AdiZ or its vector control was anaerobically cultured at pH 5.5, where the endogenous AdiZ expression was maximal (Figure 1F). Ectopic AdiZ was pulse-expressed by adding arabinose, and total RNA was extracted after 10 min for transcriptomic analysis. RNA-seq revealed significant changes in transcript levels, with 62 mRNAs showing decreased expression and 12 mRNAs showing increased expression in the AdiZ-expressing strain compared to the control strain (|Log_2_(Fold-Change)| > 1, adjusted p-value < 0.05; Figures 3A and 3B; Table S1).

**Figure 3.**
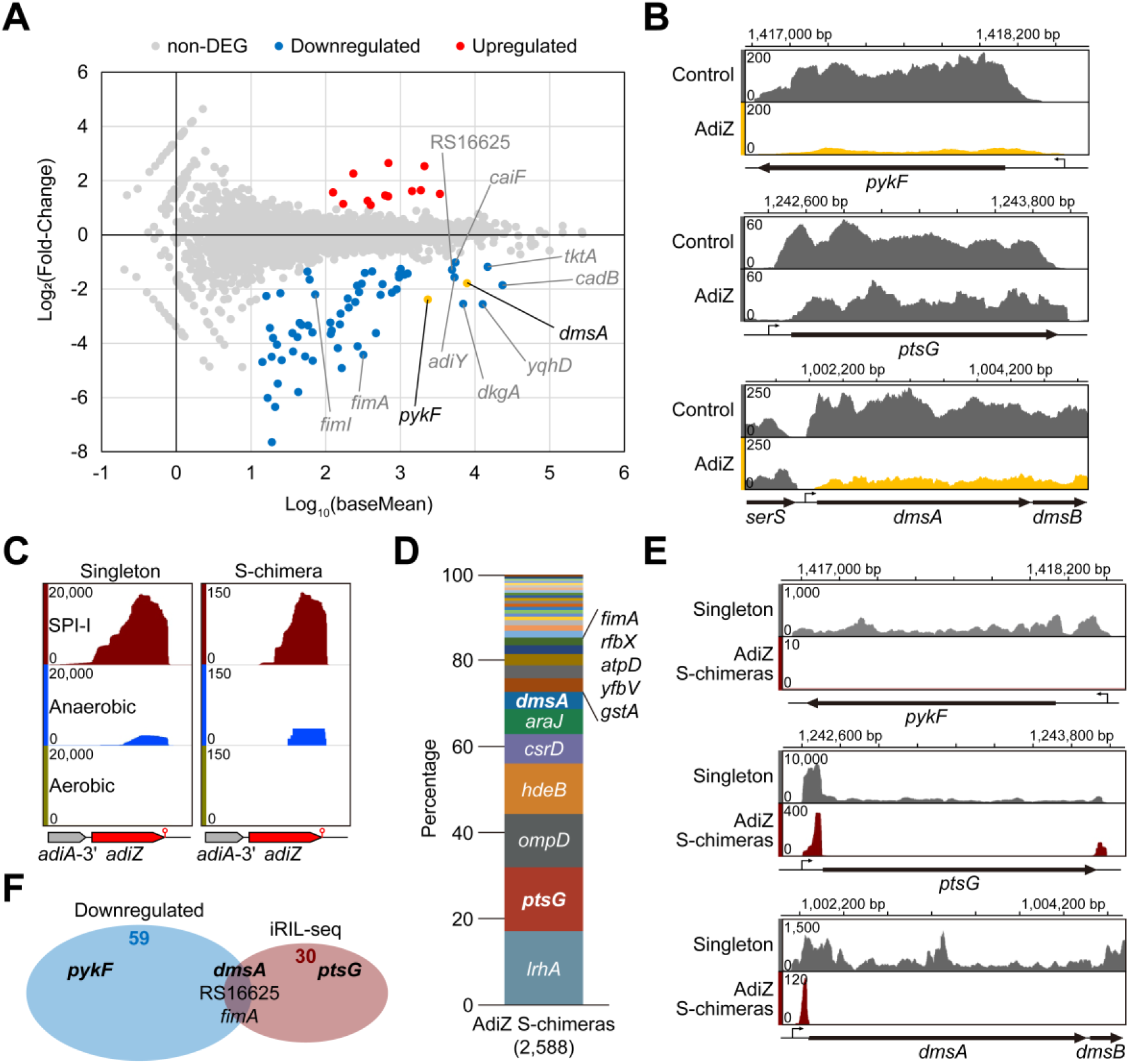
Target RNA candidates of AdiZ revealed by RNA-seq-based approaches. (A) Transcriptomic changes upon AdiZ pulse-expression. *Salmonella* Δ*adiZ* strains harboring pBAD-AdiZ or its vector control (pKP8-35) were grown anaerobically in LB (pH 5.5). At late exponential phase, 0.2% L-arabinose was added to the cultures, followed by a 10-min incubation. Differentially expressed genes (DEGs; adjusted p-value < 0.05, n = 2) are shown: blue for downregulated (Log_2_(Fold-Change) < -1) and red for upregulated (Log_2_(Fold-Change) > 1). Note that AdiZ is not represented in this dataset due to the use of a column-based mRNA enrichment protocol during RNA extraction and an annotation pipeline that excluded sRNAs. (B) Genome browser screenshots showing decreased mRNA levels of *pykF* and *dmsA* upon AdiZ pulse-expression. (C) iRIL-seq detected AdiZ under SPI-I and anaerobic conditions. Genome browser screenshots showing the genomic locations of *adiZ* covered by singleton or significant chimeric reads (S-chimera, *p* < 0.05, one-sided Fisher’s exact test). (D) Relative abundance of chimeric RNAs with AdiZ detected by iRIL-seq. Percentages were calculated as the number of fragments corresponding to each RNA divided by the total number of chimeric fragments with AdiZ. (E) Genome browser screenshots showing singleton and chimeric reads with AdiZ mapped to *ptsG* and *dmsA*. (F) Venn diagram showing the overlap between potential AdiZ targets identified by RNA-seq following pulse-expression and iRIL-seq.

Next, to identify transcripts that directly interact with AdiZ, we employed iRIL-seq analysis (intracellular <u>R</u>NA interaction by ligation and <u>seq</u>uencing) to profile RNA-RNA interactome (31). By pulse-expressing T4 RNA ligase 1 from a P_BAD_ promoter, iRIL-seq enables *in vivo* proximity ligation of sRNAs with their interaction partners in living bacterial cells. Hfq-bound ligation products (RNA chimeras) are subsequently enriched via Hfq-coIP, followed by RNA-seq analysis. Here, AdiZ reads were markedly enriched as both chimeras and non-chimeric singletons under anaerobic and *Salmonella* pathogenicity island (SPI)-1-inducing conditions (hypoxic and high-salt) (40), compared to the aerobic condition (Figure 3C). Notably, iRIL-seq under SPI-1 conditions delivered the highest number of AdiZ-containing chimeric reads, identifying a total of 33 RNAs interacting with AdiZ (Figure 3D, Table S2). For *ptsG* and *dmsA* mRNAs, chimeric reads were predominantly mapped to their 5′UTRs (Figure 3E). Only three RNAs (*dmsA*, SL1344_RS16625, and *fimA*) were identified as candidate targets by both RNA-seq-based approaches (Figure 3F).

### AdiZ represses *pykF*, *ptsG*, and *dmsA* via direct base-pairing

To assess whether AdiZ interacts with and regulates the candidate target RNAs identified by the two RNA-seq-based approaches as described above, we conducted *in silico* base-pairing predictions using IntaRNA (41), followed by translational reporter assays using the established two-plasmid system (42). For the IntaRNA analysis, candidate sequences were selected from transcription start sites retrieved from SalCom v1.0 (https://bioinf.gen.tcd.ie/cgi-bin/salcom.pl?_HL (14)), or -200 nt for intraoperonic candidates, to +200 nt relative to the start codons. Sequences with strong base-pairing potential (hybridization energy < -10 kcal/mol) to the main body of AdiZ were fused to superfolder GFP (sfGFP) derived from pXG-10sf and pXG-30sf plasmids, which are suitable for analyzing standalone and intraoperonic target mRNAs, respectively (42). An *E. coli* Δ*adiZ* mutant harboring a combination of the sfGFP-reporter plasmid and either an AdiZ-expressing plasmid or its vector control was cultured, and sfGFP fluorescence was measured. Relative fluorescence units (RFU) normalized to OD_600_ revealed that *pykF*, *ptsG*, and *dmsA* were significantly downregulated by AdiZ (≤ 50% relative to the vector control), but the other 18 candidates were not (Figures 4A and S2).

**Figure 4.**
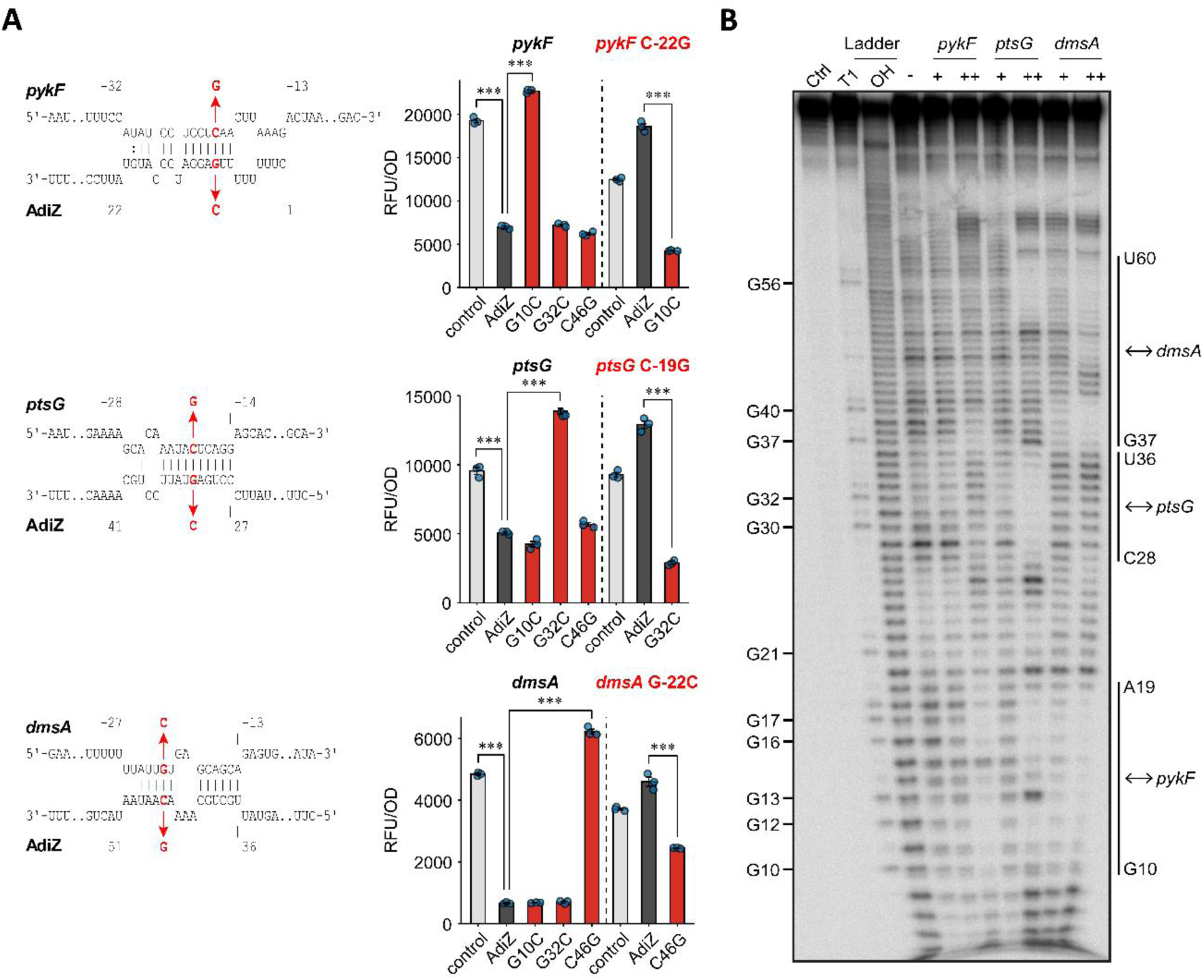
AdiZ represses *pykF*, *ptsG*, and *dmsA* expression via direct base-pairing. (A) (Left) Base-pairing interactions predicted by the IntaRNA program (41). Numbers above and below the nucleotide sequences indicate the position relative to the start codon of the mRNA and the cleavage site (i.e., 5′end) of AdiZ, respectively. (Right) sfGFP-reporter assay in *E*. *coli* Δ*adiZ* strains harboring a combination of the sfGFP-reporter plasmid and either pJV300 (vector control), pPL-AdiZ, or its derivatives expressing AdiZ point mutants (G10C, G32C, or C46G). RFU was calculated by subtracting the autofluorescence of the same strains without sfGFP-reporter plasmids and normalized to OD_600_ value. Data are presented as mean ± standard error from three independent experiments (n = 3). Statistical significance was assessed using one-way ANOVA followed by two-tailed Student’s t-test with Bonferroni correction (***p < 0.001). (B) In-line probing assay for 0.2 pmol radiolabeled AdiZ incubated with increasing concentrations of cold target mRNAs: *pykF*, *ptsG*, or *dmsA* (“+”: 0.2 pmol, “++”: 2 pmol). The “ctrl” lane represents untreated labeled RNAs, the “T1” lane represents RNase T1-digested RNAs for a G-ladder, and the “OH” lane represents alkaline-digested RNAs. Vertical labelled lines indicate protected regions corresponding to the base-pairing interactions validated in (A).

*pykF* encodes pyruvate kinase I, a key enzyme catalyzing the last step of glycolysis (33), while *ptsG* encodes the enzyme IIBC component of the glucose-specific phosphotransferase system, which mediates glucose uptake (34). *dmsA* encodes the dimethyl sulfoxide (DMSO) reductase subunit A, an essential component for DMSO respiration under anaerobic conditions (43). While *dmsA* was predicted to be an AdiZ target by both approaches, *pykF* and *ptsG* were identified as target candidates in either the pulse-expression or iRIL-seq approach (Figure 3F), illustrating the added value of orthogonal sRNA target screens.

To validate direct base-pairing between AdiZ and its target mRNAs, point mutations were introduced into both the AdiZ-expressing plasmid and the sfGFP-reporter plasmids. The G10C, G32C, and C46G mutations in AdiZ abrogated repression of *pykF*, *ptsG*, and *dmsA*, respectively, whereas mutations outside the base-pairing regions did not affect the regulation (e.g., AdiZ mutants G32C and C46G repressed PykF levels comparable to wild-type AdiZ) (Figure 4A). Conversely, complementary mutations in the target mRNAs (C-22G in *pykF*, C-19G in *ptsG*, and G-22C in *dmsA*) rather reversed the response to the wild-type AdiZ, but restored repression when combined with the corresponding AdiZ mutants. We hypothesize that AdiZ utilizes three distinct seed sequences to post-transcriptionally repress *pykF*, *ptsG*, and *dmsA* through direct base-pairing with their 5′UTRs.

To further identify the exact base-pairing regions, we probed the structure of AdiZ in association with *in vitro*-transcribed mRNA fragments (Figure 4B). A ten-fold excess of *pykF*, *ptsG*, and *dmsA* mRNAs suppressed cleavage at the AdiZ regions G10-A19, C28-U36, and G37-U60, respectively, consistent with the *in silico* predictions (Figure 4A). Binding to the *dmsA* mRNA also protected the A11-C18 region, which is complementary to the G55-U64 and forms the stem-loop in AdiZ (Figure 1E). Notably, even equimolar amounts of *dmsA* mRNA produced a similar protection pattern to that observed with a ten-fold excess. These results validate the predicted base-pair interactions and suggest that the *dmsA* mRNA may bind to AdiZ with higher affinity than *pykF* and *ptsG*.

### Physiological effects of AdiZ on the expression of its target proteins

To examine the regulatory effects of AdiZ on the endogenous protein levels of its targets in *Salmonella*, a 3x-FLAG tag was introduced at their C-termini, and protein levels were analyzed by Western blot. Under the anaerobic condition in LB medium supplemented with MES (pH 5.5), plasmid-driven overexpression of AdiZ in the Δ*adiZ* background strongly repressed all targets to <20% of the levels observed with the vector control (Figure 5A). The G10C, G32C, and C46G mutations in AdiZ abrogated the repression of *pykF*, *ptsG*, and *dmsA*, respectively, while mutations outside the base-pairing regions had no effect (Figure 5A), consistent with the results of the sfGFP-reporter assay (Figure 4A). DmsA contains a 45-amino-acid twin-arginine leader peptide, which functions as a signal for membrane targeting and is proposed to be proteolytically cleaved immediately after this sequence (43, 44). Two bands corresponding to DmsA-3x-FLAG were observed only in the vector control and the *adiZ*-C46G mutant: the upper band represents the precursor form, while the lower band corresponds to the mature form (Figure 5A). Since AdiZ overexpression decreased the levels of both forms, AdiZ likely represses *dmsA* translation rather than its post-translational processing, a conclusion supported by its interaction with the 5′UTR of *dmsA* mRNA (Figures 3E and 4A).

**Figure 5.**
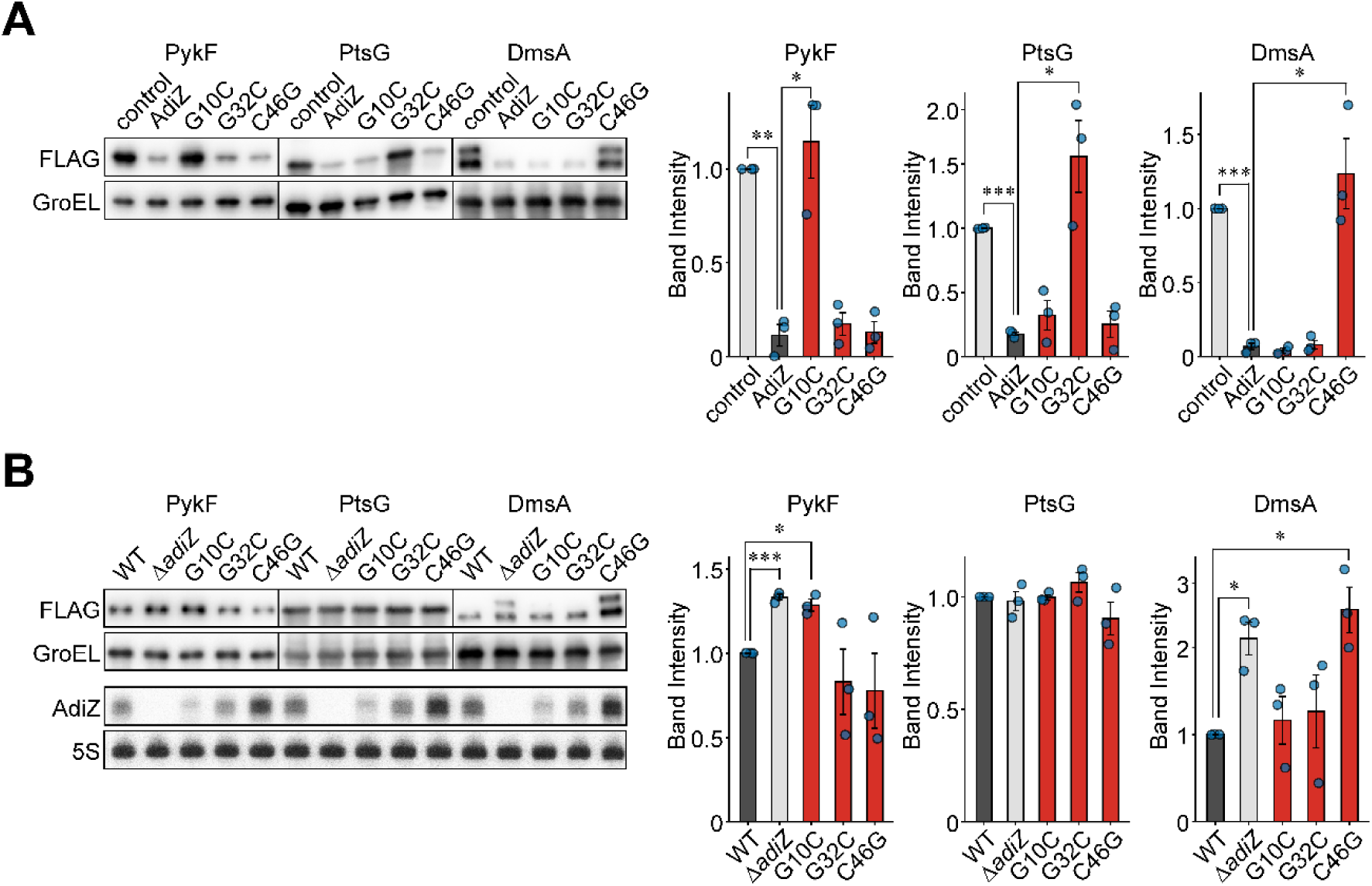
Western blot analysis of endogenous protein levels of AdiZ targets. (A) *Salmonella* Δ*adiZ* mutants with C-terminal 3x-FLAG fusions to PykF, PtsG, or DmsA were transformed with pJV300 (vector control), pPL-AdiZ, or its derivatives expressing AdiZ point mutants (G10C, G32C, or C46G). Cells were cultured anaerobically at pH 5.5 until the late exponential phase. (B) Wild-type *Salmonella*, the Δ*adiZ* mutant, and the chromosomal point mutants (*adiZ*-G10C, G32C, or C46G) with C-terminal 3x-FLAG fusions to PykF, PtsG, or DmsA were cultured as in (A). RNA extracted under the same conditions was analyzed by northern blot. AdiZ was detected with MMO-2143 and 5S rRNA detected with MMO-1056 served as a loading control. Western blot images are representative of three independent experiments. Band intensities of PykF, PtsG, and DmsA were normalized to GroEL and are presented as mean ± standard error (n = 3). Statistical significance was assessed using one-way ANOVA followed by two-tailed Student’s t-test with Bonferroni correction (*p < 0.05, **p < 0.01, ***p < 0.001).

We next examined the effect of endogenous AdiZ on target expression using the Δ*adiZ* mutant and the chromosomal point mutants. Under the same conditions, PykF levels in the Δ*adiZ* mutant and the G10C mutant were significantly increased to 1.3-fold compared to those in the wild type (Figure 5B, upper panels). For DmsA, in addition to the increased levels of the mature form, precursor forms were observed exclusively in the Δ*adiZ* mutant and the C46G mutant. Since DmsA predominantly exists in its mature form in the wild type, we suggest that AdiZ tightly regulates the translation of DmsA to keep pace with its processing. Northern blot analysis revealed that the levels of the AdiZ G10C mutant were lower, while those of the C46G mutant were higher compared to the wild type (Figure 5B, lower panels). Nonetheless, these differences in AdiZ abundance did not affect the extent of regulation since AdiZ-G10C still repressed DmsA expression. In contrast to PykF and DmsA, the PtsG levels were not significantly affected by any of the *adiZ* mutations under this condition.

The expression of *ptsG* is tightly regulated in response to glucose availability. Glucose represses *ptsG* through two mechanisms: by reducing intracellular cAMP levels, which abolishes cAMP-CRP complex-dependent activation (45), and by inducing SgrS sRNA, which downregulates *ptsG* via translation inhibition and mRNA degradation (46, 47). Consistent with this, when cells were anaerobically cultured in LB containing 0.4% glucose at pH 5.5, PtsG levels dropped markedly to ∼20% of those observed in the absence of glucose (Figure S3). Under this condition, although the level of AdiZ remains constant, PtsG levels in the Δ*adiZ* mutant and the G32C point mutant increased to ∼1.5-fold compared to those in the wild type. In contrast, PykF levels remained unaffected in all the *adiZ* mutants under this condition (Figure S3). Nonetheless, DmsA levels increased in the Δ*adiZ* mutant and the C46G mutant regardless of glucose supplementation (Figures 5B, upper panels and S3), reflecting the higher affinity of the *dmsA* mRNA to AdiZ (Figure 4B).

### AdiZ attenuates DMSO respiration coupled with glycolysis

To uncover the physiological significance of the post-transcriptional regulation by AdiZ, we screened for phenotypes of a *Salmonella* Δ*adiZ* mutant, in which the *adiA* 3′UTR upstream of the intrinsic terminator is replaced with FRT sequence. We first examined whether the deletion of *adiZ* affects the expression or activity of AdiA protein, which catalyzes arginine decarboxylation coupled with H^+^ consumption. Under the anaerobic condition in LB medium supplemented with MES (pH 5.5), the deletion of *adiZ* alone did not affect the arginine-dependent decarboxylation, while it was markedly reduced in both Δ*adiA* and Δ*adiAZ* mutants (Figure 6A). This result indicates that the expression of AdiA protein is not altered by deleting *adiZ*.

**Figure 6.**
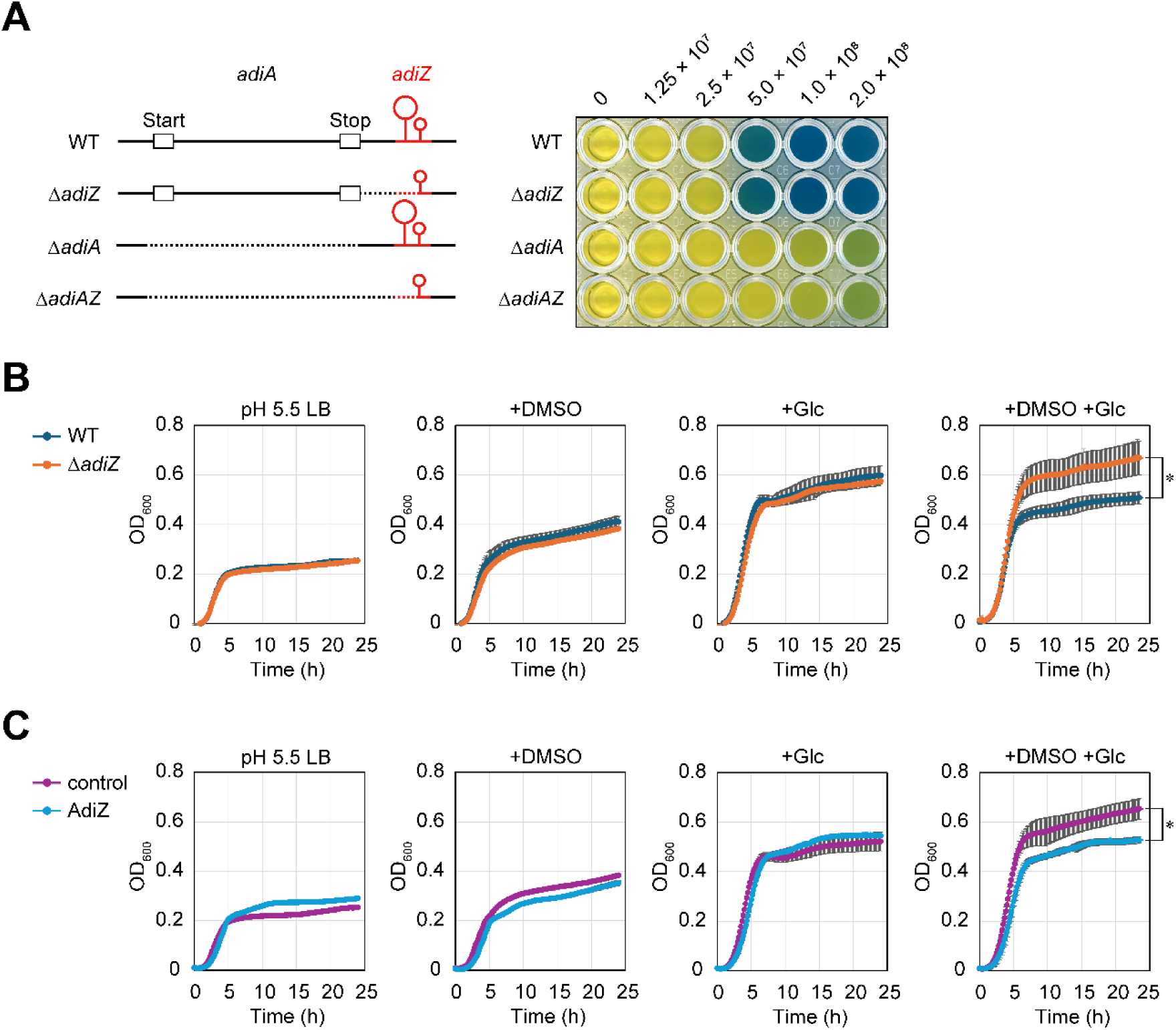
AdiZ represses *Salmonella* growth under acidic and anaerobic conditions in the presence of DMSO and glucose. (A) Deletion of *adiZ* does not affect AdiA expression. Left: schematic representation of the *adiA*- *adiZ* locus and individual deletion mutants, with deleted regions indicated by dashed lines. The intrinsic terminator remains intact in all strains. Right: *Salmonella* cells were cultured anaerobically in acidic LB (pH 5.5), washed with 0.85% NaCl, and aliquots (1.25, 2.5, 5.0, 10, or 20 × 10^7^ cells) were transferred to the arginine decarboxylase assay reagent (pH 3.4) containing bromocresol green as a pH indicator. Arginine decarboxylase raises the reagent pH, resulting in a color change from yellow to blue. (B) Biological triplicates of wild-type *Salmonella* and the Δ*adiZ* mutant were cultured anaerobically in LB (pH 5.5) in the absence or presence of 1% DMSO, 0.4% glucose, or both. Growth was continuously monitored by measuring OD_600_ every 10 min for 24 h. (C) *Salmonella* Δ*adiZ* mutants harboring either pJV300 (vector control) or pP_L_-AdiZ were cultured as in (B). Data are presented as mean ± standard error (n = 3) and OD_600_ values at 24 h were statistically analyzed using the two-tailed Student’s t-test (*p < 0.05) in (B) and (C).

Given that PykF and PtsG play key roles in glucose metabolism (33, 34) and DmsA is required for anaerobic respiration using DMSO as the terminal electron acceptor (43), we hypothesized that AdiZ might influence DMSO respiration-dependent growth of *Salmonella*. To test this, wild-type *Salmonella* and the Δ*adiZ* mutant were cultured anaerobically in LB supplemented with MES (pH 5.5) in the presence or absence of 1% DMSO, 0.4% glucose, or both for 24 h. The final cell density (OD_600_ value) of the Δ*adiZ* mutant was 1.3-fold higher than that of the wild type in the presence of both DMSO and glucose (Figure 6B). In addition, AdiZ overexpression in the Δ*adiZ* mutant reduced the final OD_600_ to ∼80% of that observed in the vector control under the same condition (Figure 6C). No significant differences were observed under conditions lacking either DMSO or glucose. These results suggest that AdiZ attenuates a series of DMSO respiration reactions, where electrons are transferred from NADH generated through glucose metabolism, thereby modulating metabolic flux under anaerobic, glucose-rich conditions.

### AdiZ contributes to *Salmonella* infection within macrophages

Recent studies indicate that both glycolytic flux and DmsA-dependent respiration are crucial for *Salmonella* survival within macrophages, where the bacterium faces a hypoxic (∼1% O_2_) and acidic (pH <5) environment (5, 35). To counteract the dissipation of ΔpH caused by reactive oxygen species (ROS) from phagocyte NADPH oxidase, *Salmonella* shifts its metabolic flux from oxidative phosphorylation to glycolysis (48). The metabolic reprogramming within macrophages is also supported by transcriptional profiling using a comprehensive reporter library (49). Additionally, phagocyte NADPH oxidase generates methionine sulfoxide (MetSO), which serves as a biologically relevant terminal electron acceptor, and DmsA-mediated respiration using MetSO further supports redox balance and contributes to cytoplasmic alkalinization in *Salmonella* (35). Based on these recent findings, we hypothesize that AdiZ, by coordinating the necessary metabolic adaptations, could play a pivotal role in *Salmonella* survival within macrophages.

To test this hypothesis, we first examined whether the *adiAZ* locus is expressed within macrophages. A transcriptional reporter plasmid was constructed by fusing the *adiA* promoter to the mCherry reporter gene. Expression of mCherry was induced under acidic conditions (Figure S4), consistent with the *adiA* mRNA expression pattern observed in Figure S1. To assess *adiA* promoter activity within host cells, sfGFP-expressing *Salmonella* harboring either the P*_adiA_*-mCherry reporter or the empty vector control were added to cultures of mouse macrophage-like RAW264.7 cells, and the fluorescence was analyzed by microscopy up to 20 hours post infection (h.p.i.). As expected, no mCherry signal was detected at any time point for the empty vector control (Figure S5). In the case of the P*_adiA_* reporter, a weak mCherry signal was detectable at 1 h.p.i., and it became stronger at 3 and 5 h.p.i. (Figures 7A and 7B). By 20 h.p.i., most intracellular *Salmonella* exhibited mCherry fluorescence, with a median of 89.4%. These results indicate that *adiAZ* expression is activated following *Salmonella* entry into macrophages.

**Figure 7.**
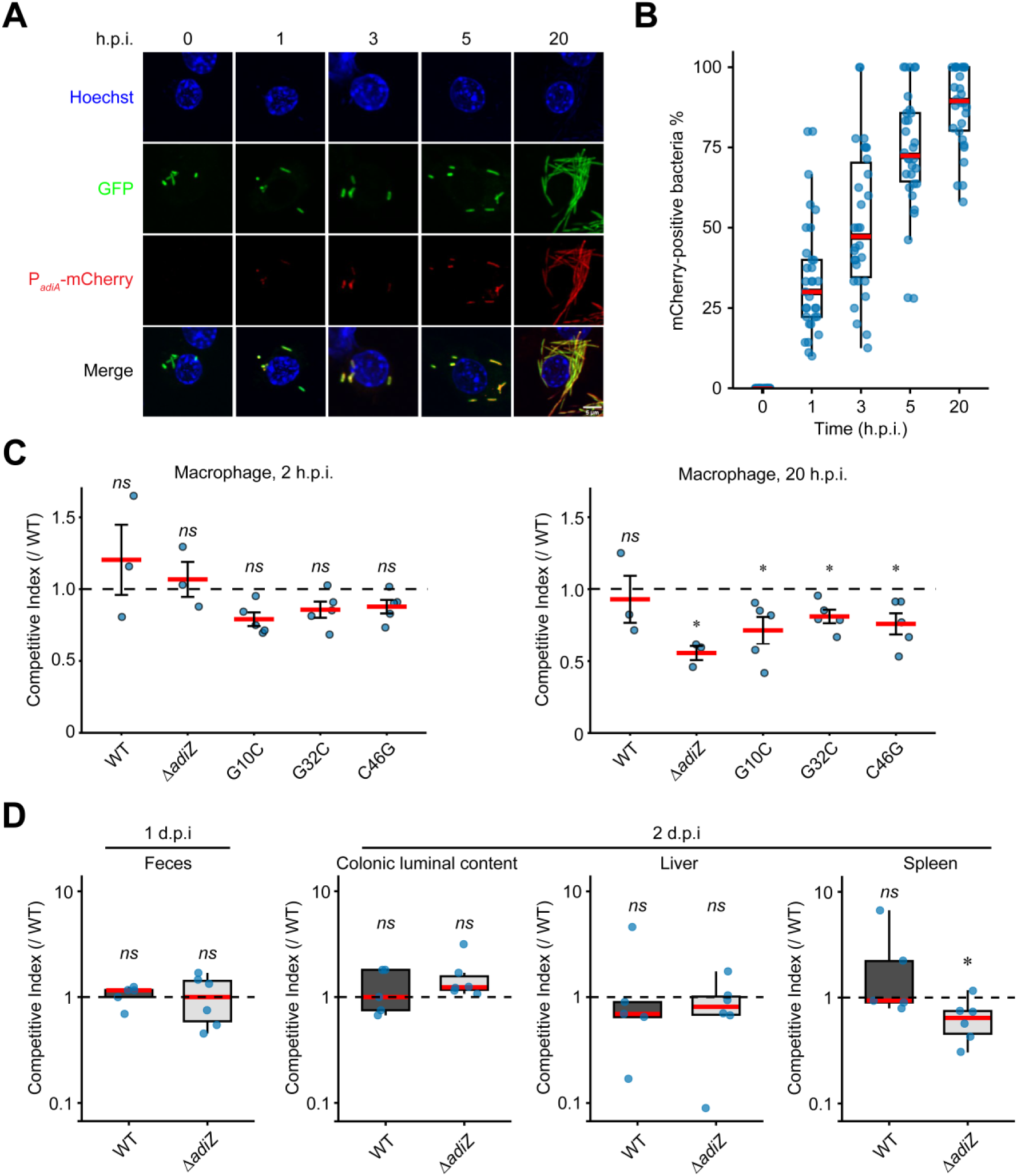
AdiZ promotes *Salmonella* infection *in vitro* and *in vivo*. (A) Activation of the *adiA* promoter following *Salmonella* entry into macrophages. An sfGFP-expressing *Salmonella* strain harboring the P*_adiA_*-mCherry reporter plasmid (pZE12-P*_adiA_*-*mcherry*) was added to cultures of mouse macrophage-like RAW264.7 cells. At the indicated time points, cells were fixed, stained with Hoechst, and analyzed by fluorescence microscopy. (B) Time-dependent proportion of *Salmonella* cells expressing mCherry within macrophages. At 0, 1, 3, 5, and 20 h.p.i., the percentage was calculated by dividing the number of mCherry-positive cells by the total number of intracellular *Salmonella*. In each of two independent experiments (n = 2), at least fifteen host cells were analyzed per time point. Due to variation in bacterial load, the number of *Salmonella* cells analyzed varied: ∼80-160 until 5 h.p.i., and ∼500 at 20 h.p.i. Red bars indicate the median with interquartile ranges. (C) Competitive infection assay between wild-type *Salmonella* and *adiZ* mutants within macrophages. *Salmonella* cells opsonized with mouse serum were added to cultures of mouse macrophage-like RAW264.7 cells. Fitness was assessed 2 and 20 h.p.i. by calculating the CI between strains carrying different fluorescence markers (sfGFP *adiZ*-WT vs. mSc *adiZ*-WT, deletion, G10C, G32C, or C46G) within host cells. The dashed line represents CI = 1. Red bars represent the mean CI with SEM (n = 3 or 5). (D) Competitive infection assay between wild-type *Salmonella* and the Δ*adiZ* mutant in mouse. *Salmonella* cells were orally administered to streptomycin-pretreated 8-9-week-old female C57BL/6J mice. Fecal samples were collected 1 d.p.i., and colonic luminal contents, livers, and spleens were harvested 2 d.p.i. Bacterial fitness was assessed by calculating the CI between strains carrying different fluorescence markers (sfGFP *adiZ*-WT vs. mSc *adiZ*-WT or Δ*adiZ*). The dashed line represents CI = 1. Red bars represent the median CI with interquartile ranges (n = 5 or 6). Statistical significance was determined using the two-tailed Student’s t-test in (C) and Mann–Whitney U test in (D). *p < 0.05, “*ns*” indicates no significant difference.

Next, we evaluated the fitness of the Δ*adiZ* mutant within macrophages by conducting competitive infection assays in an *in vitro* infection model. Mixed suspensions of *Salmonella* strains carrying a functional *hisG* (*hisG*^+^) and a fluorescence marker (sfGFP or mScarlet-I; mSc) were added to cultures of RAW264.7 cells, following opsonization with mouse serum. The competitive index (CI) was determined at 2 and 20 h.p.i. based on colony-forming units (CFUs). As an important control, competition between the wild-type strains (WT sfGFP vs. WT mSc) showed no significant fitness differences at either time point (Figure 7C). Although Δ*adiZ* CFUs did not significantly differ from wild-type CFUs at 2 h.p.i. inside the macrophages, the intramacrophage CFU counts were significantly reduced at 20 h.p.i., with a mean CI value of 0.56 (Figure 7C). No significant differences were observed in the supernatants at either time point (Figure S6). Furthermore, fluorescence marker swapping (WT mSc vs. Δ*adiZ* sfGFP) consistently reproduced the lower fitness of the Δ*adiZ* mutant (CI = 0.50, Figure S7). These results reveal the positive effect of AdiZ-mediated regulations in promoting *Salmonella* survival within macrophages.

To pinpoint which AdiZ-targeted gene(s) is (are) responsible for the above phenotype, we conducted competition assays using the *adiZ* G10C, G32C, and C46G mutant strains against the wild-type strain. Each of the three mutants exhibited a modest but significant fitness defect, with mean CI values of 0.71, 0.81, and 0.76, respectively (Figure 7C), suggesting additive effects of the disabled regulation of all three target genes. In contrast, competition between each single point mutant and the Δ*adiZ* mutant revealed no significant differences at either time point in the cells or supernatant (Figure S8). These findings suggest that AdiZ-mediated regulation of *pykF*, *ptsG*, and *dmsA* additively contributes to *Salmonella* survival within macrophages.

Lastly, we assessed the fitness of the Δ*adiZ* mutant in mice. Mixed suspensions of two strains were orally administered to 8-9-week-old female mice. Fecal samples were collected at 1 day post infection (d.p.i.) and colonic luminal contents, liver, and spleen samples were harvested at 2 d.p.i., homogenized, and plated to determine CI values based on CFU counts. As observed *in vitro*, competition between the wild-type strains (WT sfGFP vs. WT mSc) showed no significant fitness differences in any samples (Figure 7D). In contrast, the Δ*adiZ* mutant exhibited a slightly but significantly reduced fitness in the spleen compared to the wild type, with a median CI of 0.65 (Figure 7D). Together, these findings suggest that AdiZ promotes *Salmonella* survival within macrophages by fine-tuning metabolic reprogramming, thereby facilitating its establishment in the host environment (Figure 8).

**Figure 8.**
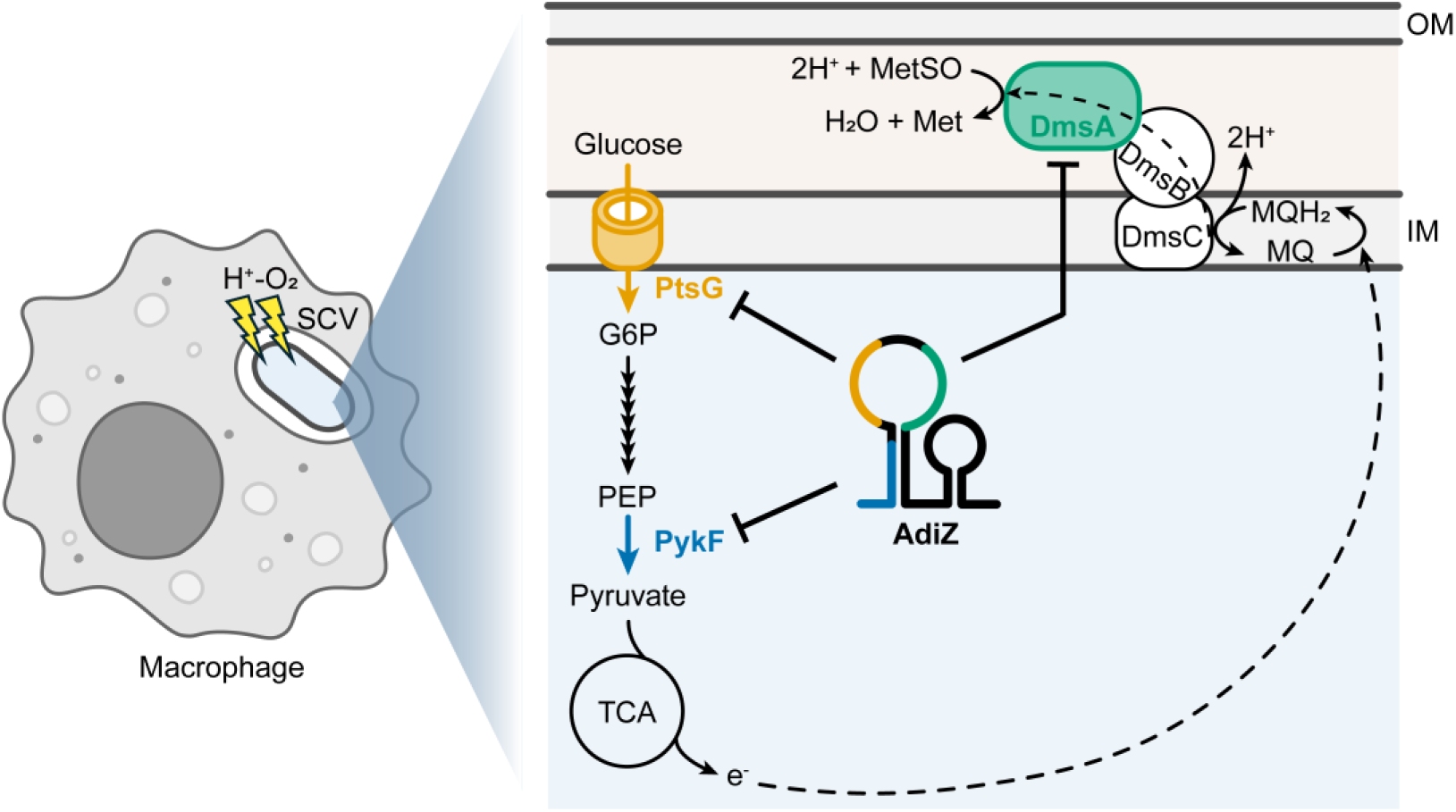
Model illustrating AdiZ-mediated fine-tuning of metabolic reprogramming in *Salmonella* within macrophages. Upon entry into macrophages, *Salmonella* encounters an acidic and hypoxic environment within the SCV, which induces the expression of AdiZ sRNA (the biogenesis mechanism is depicted in Figure 2D). AdiZ fine-tunes the reprogramming of two key metabolic pathways that support intracellular adaptation: glycolysis and MetSO-dependent anaerobic respiration, by directly targeting the *pykF*, *ptsG*, and *dmsA* mRNAs via three distinct seed regions, shown in blue, orange, and green, respectively. The flow of electrons (e^−^), generated through glycolysis and the TCA cycle and transferred to the DmsABC complex, is indicated by a dotted arrow. This coordinated modulation of metabolic gene expression enhances *Salmonella* adaptation to, and survival within, the intracellular niche. SCV, *Salmonella*-containing vacuole; OM, outer membrane; IM, inner membrane; MetSO, methionine sulfoxide; G6P, glucose-6-phosphate; PEP, phosphoenolpyruvate; TCA, tricarboxylic acid cycle; MQ, menaquinone.

## DISCUSSION

Bacterial acid resistance has been extensively studied since the 1990s due to its potential relevance to pathogenicity by enabling survival in severe acidic environments within the host. In the facultative intracellular pathogen *Salmonella*, the arginine decarboxylase encoded by the *adi* locus has been identified as the most effective system for resisting extremely acidic conditions, such as those found in the stomach (9). In this study, we extend the physiological significance of the *adi* locus beyond its classic role in acid resistance by characterizing the *adiA* 3′UTR-derived sRNA AdiZ. AdiZ is transcribed as part of the *adiA* mRNA under acidic and anaerobic conditions, cleaved by RNase E, and stabilized by Hfq (Figure 2D).

One major challenge in the characterization of AdiZ is its absence under standard culture conditions, such as aerobiosis and neutral pH. Recent studies exploring the global RNA-RNA interactome in *Salmonella* and *E. coli* under normoxic conditions failed to capture significant reads of AdiZ (31, 50–55). However, to fully uncover the infectious strategies of pathogenic bacteria, it is crucial to investigate sRNAs expressed under specific conditions especially relevant to host niches, as these regulatory RNAs could enable rapid adaptation to the dynamic environments within the host (14, 56, 57).

Our two RNA-seq-based approaches, both conducted under anaerobic conditions, successfully provided a comprehensive list of potential AdiZ target RNAs (Figure 3). Following *in silico* prediction, reporter assay, structure probing, and western blot collectively identified *pykF*, *ptsG*, and *dmsA* mRNAs as direct AdiZ targets (Figures 4 and 5). Among the three validated targets, *dmsA* was identified in both RNA-seq datasets (Figure 3F) and binds AdiZ with the highest affinity (Figure 4B), supporting its designation as the primary target of AdiZ. Moreover, the deletion or C46G mutation of chromosomal *adiZ* strikingly increased the levels of the premature form of DmsA (Figure 5B), suggesting that AdiZ inhibits the translation initiation of DmsA to coordinate the processing of leader peptide and the translocation of the mature form into the membrane (43, 44). It is noteworthy that DmsA is involved in cytoplasmic alkalinization of intracellular *Salmonella* by anaerobically respirating MetSO (35). Under anaerobic and acidic conditions, the *adiAZ* mRNA produces both the arginine decarboxylase and AdiZ sRNA, and the post-transcriptional regulation of *dmsA* balances the expression of two distinct acid resistance systems in the intracellular *Salmonella*.

Notably, however, *pykF* was not contained in chimeras with AdiZ in the iRIL-seq dataset, whereas *ptsG* was not downregulated upon AdiZ pulse-expression (Figures 3B, 3E, and 3F). The lack of detectable *pykF*-AdiZ chimeras may be attributed to the relatively high abundance of ArcZ sRNA-*pykF* chimeras under the same condition (Figure S9), which perhaps masked the detection of less abundant interactions. Conversely, *ptsG* may be regulated by AdiZ primarily at the translational level without affecting mRNA abundance, explaining its presence in the iRIL-seq data but not in the pulse-expression RNA-seq data. *ptsG* is also a well-known target of the SgrS sRNA, which promotes its rapid degradation during glucose-phosphate stress (46). Intriguingly, *Salmonella* and *Citrobacter* AdiZ harbor and utilize the complete six-nucleotide seed sequence of SgrS that forms the critical core interaction with *ptsG* mRNA (58), whereas this sequence is not conserved in *Escherichia* or *Shigella* (Figure S10). Despite targeting the same site, AdiZ and SgrS appear to exert mechanistically distinct modes of regulation: SgrS triggers mRNA degradation, while AdiZ modulates translation without altering transcript stability. AdiZ may act on the residual pool of *ptsG* transcripts that escape SgrS-mediated degradation, potentially explaining why AdiZ-dependent modulation of PtsG protein levels becomes detectable only under the glucose-replete condition (Figure S3).

### sRNAs orchestrate *Salmonella* virulence and sugar metabolism

As an intriguing finding of the present study, the AdiZ G10C and G32C mutants, which have lost their ability to regulate *pykF* and *ptsG*, respectively, exhibited significantly reduced fitness within macrophages compared to the wild type (Figure 7C). This emphasizes the importance of sRNA-mediated fine-tuning of sugar metabolism for intracellularly surviving *Salmonella*.

Recent findings provide further evidence for the regulatory interplay between *Salmonella* virulence and sugar metabolism (59). The *Salmonella*-specific sRNA PinT, for example, is a key regulator of virulence, controlling both SPI-1 and -2 by regulating key transcription factors HilA, RtsA, and SsrB, as well as the global carbon metabolism regulator CRP (60, 61). CRP suppression by PinT influences SPI-2 expression during intracellular survival (60). Furthermore, the acid-inducible sRNA RyeC represses translation of PtsI, a key component of the phosphoenolpyruvate:sugar phosphotransferase system, and promotes intracellular survival within macrophages (62). More recently, the 3′UTR-derived sRNA ManS was shown to contribute to *Salmonella* colonization in the host by coordinating sialic acid metabolism (22).

SgrS represses the expression of multiple sugar transporters including PtsG in response to sugar-phosphate stress, and specifically in *Salmonella*, also directly regulates the SPI-1 effector SopD (63), suggesting that *Salmonella* senses subcellular host environments via carbon source availability and fine-tunes effector expression. Intriguingly, SgrS also directly base-pairs with *adiY* mRNA, leading to its translational inhibition and degradation in *E. coli* (64). Our findings that AdiY strongly induces AdiZ expression and AdiZ represses *ptsG* thus suggest the existence of a coordinated sRNA-based regulatory circuit that ensures precise and robust control of *ptsG* expression in response to environmental cues.

### AdiA-dependent acid resistance in the SCV environment

Inside macrophages, *Salmonella* encounters an environment characterized by acidity and hypoxia, namely SCV. To adapt to this hostile niche, SCV acidity is sensed by *Salmonella*, which induces SPI-2 through a cascade of EnvZ/OmpR, PhoQ/PhoP, and SsrA/SsrB two-component systems (65). OmpR and PhoP activate the transcription of *ssrA* and *ssrB*, respectively (66), and SsrB directly responds to acidic pH through a conserved histidine residue to stimulate the transcription of SPI-2 genes encoding the type-III secretion system (T3SS) and its effector proteins required for intracellular proliferation (67). Interestingly, OmpR also represses the CadA/CadB acid resistance system to prevent cytoplasmic neutralization (68). How these regulators control the transcription of *adi* locus in conjunction with AdiY requires further investigation.

Arginine is one of the seven essential metabolites that support *Salmonella* colonization in mice (69) and is critical for intracellular *Salmonella* to withstand ROS and nitric oxide (NO) produced by phagocyte NADPH oxidase and inducible NO synthase (iNOS), respectively (70). In addition to *de novo* synthesis, *Salmonella* acquires host-derived arginine through two ABC transporter systems and a putative L-Arg/Orn antiporter, which limit host NO production by sequestering arginine away from the iNOS pathway (70–72). Furthermore, *Salmonella* utilizes polyamines generated through host arginine catabolism to facilitate T3SS assembly (73). Given that the L-Arg/Agm antiporter AdiC is not required for *Salmonella* virulence in mice (70), the Adi system seems to be dispensable for arginine import. Neither *adiA*, *cadA*, nor *speF* was previously reported to be essential to *Salmonella* during systemic infection (9).

However, we argue that AdiZ-mediated post-transcriptional regulation of *pykF*, *ptsG*, and *dmsA* takes place within macrophages, where *Salmonella* faces an acidic, hypoxic, and arginine-rich environment. We observed induction of the *adiAZ* promoter upon *Salmonella* entry into macrophages (Figures 7A and 7B). Moreover, the *adiZ* deletion mutant displayed reduced fitness in both macrophages and mice (Figures 7C and 7D), revealing a previously unappreciated contribution of the *adi* locus to intracellular adaptation. Based on our comprehensive target identification, we propose that AdiZ facilitates *Salmonella* infection by fine-tuning two key metabolic pathways: glycolysis and DmsA-dependent anaerobic respiration. Glucose transport via PtsG and subsequent catabolism are crucial to mitigate ROS-induced stress within macrophages (48). DmsA, the primary AdiZ target, is essential for anaerobic respiration using MetSO as a terminal electron acceptor, contributing to redox balance and cytoplasmic alkalinization (35). Notably, point mutations in three distinct AdiZ seed regions (G10C, G32C, and C46G), each impairing regulation of the specific target gene, individually reduced intracellular fitness (Figure 7C), suggesting that coordinated regulation of all three targets collectively enhances adaptation to the hostile intracellular milieu.

Our mCherry-based reporter detected activation of the *adiAZ* promoter following macrophage entry (Figures 7A and 7B). However, a recent transcriptional profiling using a comprehensive GFP-reporter library failed to detect such activation (49). One possible explanation for this discrepancy is that GFP—but not mCherry—is highly sensitive to acidic pH (74); thus, genes induced in response to SCV acidification may be missed when using GFP-based reporter systems. While RNA-seq-based studies did not identify increased expression of the *adi* locus inside murine or porcine macrophages (57, 60), early transcriptome profiling using microarray analysis did detect an upregulation of *adiY* within murine macrophages (75). Transcriptional activation of the *adi* locus may therefore depend on the timing of infection, possibly requiring both hypoxia and acidification, which are gradually established inside the SCV during maturation. Noteworthy, transposon-directed insertion-site sequencing identified the *adi* locus as crucial for systemic infection in mice and intestinal colonization in chickens, pigs, and cattle (76). Disruption of the *adiAZ* terminator significantly reduced fitness in pigs and cattle (76), which may further underscore the importance of the AdiZ-mediated regulations during infection. Further studies will be required to elucidate when AdiA functions within macrophages and how it influences host arginine and polyamine metabolism.

## EXPERIMENTAL MODEL AND SUBJECT DETAILS

### Bacterial strains and growth conditions

The derivatives of *Salmonella enterica* serovar Typhimurium strain SL1344 and *Escherichia coli* used in this study are listed in Table S3. Bacterial cells were grown at 37°C, or 30°C for temperature-sensitive mutants, in LB Miller medium either aerobically with reciprocal shaking at 180 rpm or anaerobically under 76% N_2_, 20% CO_2_, 4% H_2_ atmosphere in a Coy anaerobic chamber (COY). Where required, the culture pH was adjusted to 8.0 or 5.5 with 100 mM 3-morpholinopropanesulfonic acid (MOPS, pH 8.6) or 100 mM 2-morpholinoethanesulfonic acid (MES, pH 5.4), respectively. Media were supplemented with antibiotics when necessary at the following concentrations: 50 µg/ml ampicillin (Amp), 50 µg/ml kanamycin (Km), 12.5 µg/ml chloramphenicol (Cm), and 20 mg/ml tetracycline (Tet).

### Animal studies

Animal experiment procedures were approved by the University of Tsukuba animal experiment committee. 8-week-old female C57BL/6J mice were purchased from CLEA Japan and housed in groups of 5 or 6 in ventilated cages. Mice were orally administered 20 mg of streptomycin 24 h before *Salmonella* inoculation.

*Salmonella* cells grown aerobically in LB (pH 7.0) for 21 h were harvested by centrifugation (10 min at 3,200 *g*, 20°C), washed twice with PBS, and diluted in PBS to a concentration of 10^7^ CFU/ml. Bacterial suspensions of the two strains were then mixed and 100 μl (5×10^5^ CFU of each strain) was orally administered to the mice. Fresh fecal samples were collected at 24 h.p.i. Mice were euthanized 48 h.p.i. and colonic luminal contents, livers, and spleens were harvested. The samples were homogenized, serially diluted in PBS, and plated onto LB agar containing Cm. The plates were incubated at 37°C overnight. Fluorescence signals from sfGFP or mSc within the colonies were detected using a trans illuminator (BIO CRAFT) to determine the CFU of each strain.

## METHOD DETAILS

### Plasmid construction

The plasmids and oligonucleotides used in this study are listed in Tables S4 and S5, respectively. For cloning of the *adiA*-*adiZ* and *adiZ* regions, DNA fragments were amplified using primer pairs MMO-1496/MMO-0544 and MMO-0543/MMO-0544, respectively. The amplified fragments were digested with XbaI and ligated into the vectors pKP8-35 (for arabinose-inducible expression (77)) or pJV300 (for constitutive expression (78)). The plasmid-borne *adiZ* was mutated by inverse PCR using overlapping primer pairs: MMO-1933/MMO-1934 for G10C, MMO-2095/MMO-2096 for G32C, and MMO-1807/MMO-1808 for C46G, followed by DpnI digestion. Translational fusion plasmids based on pXG10-sf and pXG30-sf were constructed as described previously (42, 79). Single-nucleotide mutations were introduced by inverse PCR using overlapping primer pairs: MMO-1935/MMO-1936 for *pykF* G10C, MMO-2093/MMO-2094 for *ptsG* G32C, and MMO-1815/MMO-1816 for *dmsA* C46G, followed by DpnI digestion. pYC963 was generated by amplifying the backbone from pJV300 using primer pair YCO-3283/YCO-3284, and the mCherry insert from pCWU6-mCherry using YCO-3285/YCO-3286. The two fragments were assembled using the Ready-to-Use Seamless Cloning Kit (Sangon Biotech). To construct pYC964, the plasmid backbone was amplified from pYC963 using YCO-3367/YCO-3368, and the *adiA* promoter region was amplified using YCO-3369/YCO-3370. The resulting fragments were similarly fused using the Ready-to-Use Seamless Cloning Kit. pXG-10-*tnaC*_eco_5′UTR-sfGFP was constructed by inverse PCR with MMO-1937/MMO-1987 using pXG-10sf-*tnaC*_eco_ as a template, followed by self-ligation of the PCR products. For pXG-10-*tnaC*_eco_5′UTR-mSc, the sfGFP sequence in pXG-10sf-*tnaC*_eco_ was removed by inverse PCR with MMO-1982/MMO-2000. The mSc sequence (80), codon-optimized for efficient expression in bacteria, was synthesized and amplified by PCR with MMO-1998/MMO-1999. Both fragments were digested with NheI and XbaI, then ligated.

### Strain construction

Deletion mutants were constructed using pKD46 for the λ Red recombination (81) and pKD13 as a template for amplifying the Km resistance cassette. The resulting Km-resistant strains were confirmed by PCR, and the mutant loci were transduced into appropriate genetic backgrounds using P22 and P1 phages for *Salmonella* and *E. coli*, respectively. The Δ*adiZ* mutants of both *Salmonella* and *E. coli* were transformed with the temperature-sensitive plasmid pCP20 expressing FLP recombinase to excise the resistance gene from the chromosome. The *rne-1* allele was transduced from strain TK40 (82) into the *E. coli* Δ*adiZ*::FRT strain using P1 phage.

Chromosomal point mutations in *adiZ* (G10C, G32C, and C46G) were introduced by scarless mutagenesis using the two-step λ Red system (83). A DNA fragment containing a Cm resistance marker and an I-SceI recognition site was amplified with primer pairs MMO-2126/MMO-2127 from the template plasmid pWRG100 and integrated into the *adiZ* region of the chromosome by λ Red recombinase expressed from pKD46. The resultant mutant was transformed with pWRG99. The mutant alleles amplified from pPL-AdiZ derivatives (Table S4) using MMO-2129/MMO-2130 for G10C or MMO-2128/MMO-2130 for G32C and C46G were integrated into the chromosome by λ Red recombinase expressed from pWRG99. Recombinants were selected on LB agar plates supplemented with Amp and 5 µg/ml anhydrotetracycline to induce I-SceI endonuclease. Successful recombinants were confirmed by Cm sensitivity, PCR, and sequencing.

For Western blot detection of PykF, PtsG, and DmsA, a 3xFLAG epitope tag was fused to the C-terminus of the corresponding proteins. Primer pairs MMO-2146/MMO-2147 and MMO-2148/MMO-2149 were used for PykF and DmsA tagging, respectively, with the template plasmid pSUB11 (84) for PCR amplification. The PCR products were integrated into the chromosome by λ Red recombinase expressed from pKD46. The resulting Km-resistant strains were confirmed by PCR, and the mutant loci were transduced into appropriate genetic backgrounds using P22 phage. For tagging PtsG, a *ptsG*-3xFLAG-Km^R^ allele was transduced from strain JVS-4141 (85) using P22 phage.

For macrophage and mouse infection assays, the *hisG* mutation causing histidine auxotrophy in the *Salmonella* SL1344 strain was restored to functionality by introducing a C206T (Pro -> Leu) reverse mutation via two-step PCR. In the first PCR step, primer pairs MMO-1943/MMO-1946 and MMO-1944/MMO-1945 were used with genomic DNA of wild-type *Salmonella* as the template. The purified products were used as templates for the second PCR step with primer pair MMO-1943/MMO-1945. The final PCR product was transformed into a *Salmonella* strain harboring pKD46 and plated onto M9 minimal medium supplemented with 0.4% glucose. The resulting strains were confirmed by sequencing, and the mutant locus was transduced into the appropriate genetic backgrounds using P22 phage.

To introduce fluorescence markers into the *Salmonella* chromosome, a DNA fragment, P_LtetO_-*tnaC*_eco_5′UTR-sfGFP-Cm^R^ or P_LtetO_-*tnaC*_eco_5′UTR-mSc-Cm^R^, was amplified by PCR using primer pairs MMO-1988/MMO-1989 and the template plasmid pXG-10-*tnaC*_eco_5′UTR-sfGFP or pXG-10-*tnaC*_eco_5′UTR-mSc (Table S4), respectively. The amplified fragment was integrated into the chromosomal *putA*-*putP* intergenic region by λ Red recombinase expressed from pKD46. The resulting Cm-resistant strains were confirmed by fluorescence and PCR, and the mutant loci were transduced into appropriate genetic backgrounds using P22 phage.

### Structure probing

DNA templates for *in vitro* transcription were amplified from wild-type *Salmonella* genomic DNA using the following primer pairs carrying a T7 promoter on the sense oligo: AWO-1787/AWO-1788 for *adiZ*, AWO-1801/AWO-1802 for *pykF*, AWO-1803/AWO-1804 for *ptsG*, and AWO-1805/AWO-1806 for *dmsA*. For each reaction, 0.4 pmol of labeled AdiZ RNA was denatured and incubated at 37°C for 15 min in the presence of RNA Structure Buffer (Thermo Fisher Scientific) and 1 µg of yeast RNA (Thermo Fisher Scientific) in a final volume of 10 µL. Lead(II)-induced cleavage was initiated by adding 2 µL of 25 nM lead(II) acetate, followed by incubation at 37°C for 90 s. Reactions were terminated by adding 12 µL of Gel Loading Buffer II (Thermo Fisher Scientific). A control reaction was performed by denaturing 1 pmol of labeled RNA at 95°C in 10 µL of water, followed by immediate cooling on ice with 10 µL of GL II RNA loading dye. To generate an alkaline hydrolysis ladder, 1 pmol of labeled RNA was incubated in Alkaline Hydrolysis Buffer (Thermo Fisher Scientific) at 95°C for 5 min. For RNase T1 digestion, 1 pmol of labeled RNA was first denatured in water at 95°C for 1 min, then incubated with RNase T1 at 37°C for 3 min. All reactions were stopped using the same procedure described above. 10 µL of each reaction was heated at 95°C for 3 min, then loaded onto a 10% polyacrylamide gel containing 7 M urea and subjected to electrophoresis at 45 W for 3 h.

### In-line probing assay

To identify nucleotide positions involved in RNA-RNA interactions, in-line probing was carried out using radiolabeled *in vitro* transcribed AdiZ RNA (0.2 pmol, 5′end-[³²P]). The RNA was incubated at room temperature for 40 h in a reaction buffer composed of 100 mM KCl, 20 mM MgCl₂, and 50 mM Tris-HCl (pH 8.3), in the presence of 0, 0.2, or 2 pmol of unlabeled target RNAs. Reactions were stopped by adding 10 µL of denaturing gel-loading buffer containing 10 M urea and 1.5 mM EDTA (pH 8.0).

To generate an RNase T1 digestion ladder, 0.4 pmol of labeled RNA was first heat-denatured at 95°C for 1 min in sequencing buffer (Ambion), followed by treatment with 0.1 U of RNase T1 at 37°C for 5 min. For the alkaline hydrolysis ladder, 0.4 pmol of labeled RNA was incubated in 9 µL of alkaline hydrolysis buffer (Ambion) at 95°C for 5 min. Both reactions were stopped with 12 µL of loading buffer II and kept on ice. RNA fragments were resolved on a 10% polyacrylamide gel containing 7 M urea, followed by electrophoresis at 45 W for 2–3 h. Gels were then dried and visualized by phosphorimager (FLA-3000 Series, Fuji).

### Northern blot

For analysis of endogenous AdiZ, *Salmonella* cells were cultured in LB (pH 7.0) either aerobically or anaerobically until the late exponential phase (OD_660_ = 0.5). The culture pH was then shifted to 8.0, 5.5, or maintained at 7.0 by adding 100 mM MOPS (pH 8.6), 100 mM MES (pH 5.4), or H_2_O, respectively. After the pH shift, cells were cultured for an additional 30 min. For *Salmonella* Δ*adiZ* strains harboring pBAD-AdiAZ or pBAD-AdiZ, 0.2% L-arabinose was added to the cultures, and cells were further incubated for 10 min. For *E. coli* Δ*adiZ* Δ*hfq* strains harboring pBAD-AdiAZ or pBAD-AdiZ, cells were cultured aerobically at 37°C until OD_660_ reached 1.0, then 0.2% L-arabinose was added, followed by a 10-min incubation. For an *E. coli* Δ*adiZ rne-1* strain harboring pBAD-AdiAZ, cells were cultured aerobically at 30°C until OD_660_ reached 0.7. The temperature was then shifted to 42°C or maintained at 30°C, and cells were cultured for an additional 30 min. Subsequently, 0.2% L-arabinose was added, followed by a 10-min incubation. Cell cultures were immediately mixed with 20% (v/v) of stop solution (95% ethanol, 5% phenol), and total RNA was isolated using the TRIzol reagent (Invitrogen).

Total RNA (5 µg) was separated on 6% polyacrylamide/7 M urea gels in 1×TBE buffer at 300 V for 2.5 h using a Biometra Eco-Maxi system (Analytik-Jena). DynaMarker RNA Low II ssRNA fragment (BioDynamics Laboratory) and/or Low Range ssRNA Ladder (NEB) served as size markers. RNA was transferred onto a Hybond N+ membrane (Cytiva) by electroblotting at 50 V for 1 h using the same device. The membrane was UV-crosslinked at 120 mJ/cm^2^, prehybridized in Rapid-Hyb buffer (Cytiva) at 42°C for 1 h, and hybridized overnight at 42°C with a [^32^P]-labeled probe (Table S5). The membrane was washed in three 15-min steps with 5×SSC/0.1% SDS, 1×SSC/0.1% SDS, and 0.5×SSC/0.1% SDS buffers at 42°C. Signals were visualized on Typhoon FLA7000 scanner (GE Healthcare).

### qRT-PCR

cDNA was synthesized using ReverTra Ace qPCR RT Master Mix with gDNA Remover (Toyobo). qRT-PCR was performed using TB Green Premix Ex Taq™ II (Takara Bio) on QuantStudio 5 Real-Time PCR System (Thermo Scientific). Each target-gene mRNA level was normalized to a reference gene transcript (*rpoB* mRNA) from the same RNA sample. Fold changes were determined using the 2^−DD*Ct*^ method (86). The sequences of the primers used are shown in Table S5.

### RNA-seq

Biological duplicates of *Salmonella* Δ*adiZ*::FRT strains harboring pKP8-35 or pBAD-AdiZ were cultured in LB (pH 7.0) anaerobically until OD_660_ reached 0.5. The culture pH was shifted to 5.5 by adding 100 mM MES (pH 5.4), and cells were cultured for an additional 30 min. Subsequently, 0.2% L-arabinose was added, and cells were further incubated for 10 min. To stabilize cellular RNA, two volumes of RNA Protect Bacterial Reagent (Qiagen) were added to one volume of the cultures. Total RNA was extracted using NucleoSpin RNA (Macherey-Nagel) according to the manufacturer’s instruction and treated with TURBO DNase (Invitrogen) at room temperature for 30 min. DNase I was then denatured and removed via phenol‒chloroform extraction, followed by ethanol precipitation. RNA quality was assessed using Bioanalyzer 2100 with the Agilent RNA 6000 Nano Kit (Agilent Technologies).

For cDNA library preparation, ribosomal RNA was depleted using the Illumina Ribo-Zero Plus rRNA Depletion Kit (Illumina). Strand-specific cDNA libraries were generated using the NEBNext Ultra II Directional RNA Library Prep Kit (Illumina) after transcript fragmentation. Libraries were sequenced with 150-bp paired-end reads on a NovaSeq 6000 system (Illumina). Raw reads were quality-checked using FastQC (ver. 0.11.7), and low-quality bases (< 20) and adapter sequences were trimmed using Trimmomatic (ver. 0.38) with the following parameters: ILLUMINACLIP:path/to/adapter.fa:2:30:10 LEADING:20 TRAILING:20 SLIDINGWINDOW:4:15 MINLEN:36. Trimmed reads were aligned to the *S*. *enterica* subsp. *enterica* serovar Typhimurium strain SL1344 reference genome (ASM21085v2) using HISAT2 (ver. 2.1.0). Details of read counts and mapping rates are shown in Table S6. The resulting .sam files were converted to .bam files using Samtools (ver. 1.9), and uniquely mapped reads were quantified with featureCounts (ver. 1.6.3). Differential expression analysis was performed with DESeq2 (ver. 1.24.0), using relative log normalization (RLE). Genes were considered differentially expressed if |Log2(Fold-Change)| > 1 and the adjusted p-value (Benjamini-Hochberg method) was < 0.05. Original sequence reads have been deposited in the DRA database under accession numbers DRR689954–DRR689957.

### iRIL-seq procedure

iRIL-seq was performed as previously described (31), with minor modifications. A *Salmonella hfq*::3xFLAG strain harboring the plasmid pBAD-*t4rnl1* (pYC582) was grown overnight in LB medium at 37°C with shaking at 220 rpm. For aerobic and anaerobic conditions, overnight cultures were diluted 1:100 into fresh LB and grown under the respective conditions. For SPI-1-inducing conditions, cultures were diluted 1:100 into LB supplemented with 0.5 M NaCl and incubated anaerobically. Two biological replicates were conducted for each condition. When cultures reached an OD_600_ of 0.3, T4 RNA ligase expression was induced by adding 0.2% L-arabinose for 30 min. Cells corresponding to 50 OD_600_ units (e.g., 100 ml at OD_600_ = 0.5) were harvested by centrifugation at 12,000 *g* for 5 min at 4°C. Pellets were washed twice with 10 ml pre-chilled PBS and stored at −80°C until further use.

Hfq co-IP was carried out as described in (87). Bacterial pellets were resuspended in 600 µl ice-cold lysis buffer (20 mM Tris-HCl pH 8.0, 150 mM KCl, 1 mM MgCl₂, 1 mM DTT, 0.05% Tween-20). Cells were lysed by bead-beating with 500 µl glass beads using a Cryolys Evolution homogenizer (Bertin Technologies) at 4°C for 10 min. Lysates were cleared by centrifugation at 17,000 *g* for 30 min at 4°C and transferred to fresh tubes. For co-IP, lysates were incubated with anti-FLAG antibody (Sigma-Aldrich)-conjugated protein G magnetic beads (Thermo Fisher Scientific) for 1 h at 4°C with gentle rotation. Beads were washed five times with 500 µl ice-cold lysis buffer and resuspended in 100 µl of the same buffer. Hfq-associated RNA was purified using RNA Clean & Concentrator columns (Sangon Biotech) and eluted in nuclease-free water.

iRIL-seq libraries were prepared using a modified RNAtag-Seq protocol through the following steps (88). Briefly, RNA was fragmented, treated with DNase I, and purified using 2.5×Agencourt RNAclean XP beads (Beckman Coulter) and 1.5×isopropanol. RNA was ligated to a 3′-barcoded adaptor and purified with 2.5×AMPure XP beads and 1.5×isopropanol. Ribosomal RNA was removed using the Ribo-off rRNA Depletion Kit (Vazyme), followed by purification with 2.5×RNAclean XP beads and 1.5×isopropanol. First-strand cDNA was synthesized with HiScript II First Strand cDNA Synthesis Kit (Vazyme). Residual RNA was hydrolyzed with 1 M NaOH, and cDNA was purified using 2.5×AMPure XP beads and 1.5×isopropanol. The cDNA was ligated with a second adaptor and purified twice with 2.5×RNAclean XP beads and 1.5×isopropanol. Libraries were PCR amplified with Illumina P5 and P7 primers using Q5 High-Fidelity DNA Polymerase (NEB), and purified with 1.5×RNAclean XP beads. Final libraries were sequenced on an Illumina NovaSeq 6000 platform with 150 bp paired-end reads.

iRIL-seq data were analyzed as described previously (31). The raw sequencing reads were processed with Cutadapt to remove adapter sequences and low-quality ends. The first 25 nt of high-quality paired-end reads were mapped to the *Salmonella* SL1344 genome (NC_016810.1) using BWA software with default parameters. The paired fragment ends were designated as two mates. Two mates of 25 nt fragments that mapped within a distance of 1,000 nt or within the same transcript were considered as singletons, whereas fragments that mapped to two different loci were defined as chimeras. Fisher’s exact test was applied to assign an odds ratio and a *p*-value to each chimera. Chimeras with a *p*-value ≤ 0.05 and an Odds Ratio ≥ 1 were defined as S-chimeras. Only S-chimeras covered by ≥ 10 chimeric reads were considered for further analysis.

### GFP fluorescence quantification

The *E. coli* MG1655 Δ*adiZ*::FRT strains harboring a combination of the sfGFP translational fusions and pJV300, pP_L_-AdiZ, or its derivatives were grown in 100 μl of LB (pH 7.0) in 96-well optical bottom black microtiter plates (Thermo Scientific). The plates were incubated at 37℃ with rotary shaking at 180 rpm and a 3-mm amplitude in Humidity Cassette using Spark plate reader (Tecan). Both OD_600_ and fluorescence (excitation at 485 nm and emission at 535 nm with dichroic mirror of 510 nm, fixed gain value of 50) were continuously measured every 10 min. At OD_600_ of 0.6, the relative fluorescence unit (RFU) was calculated by subtracting the autofluorescence of the same strains without the sfGFP-reporter plasmids, then normalized to the OD_600_ value.

### Western blot

*Salmonella* cells were cultured anaerobically in pH 5.5 LB supplemented with 100 mM MES (pH 5.4) in the absence or presence of 0.4% glucose. When OD_660_ reached 0.4, the cultures were collected by centrifugation at 3,200 *g* for 10 min at 4°C. The pellets were dissolved in 1×Laemmli Sample Buffer (Bio-Rad) to a final concentration of 0.01 OD/µl, then heated at 95°C for 5 min. For PykF-3xFLAG analysis, 0.0007 OD of whole-cell samples was separated on 10% TGX gels (Bio-Rad). For PtsG-3xFLAG and DmsA-3xFLAG analysis, 0.01 OD of each sample was separated on 10% and 7.5% gels, respectively. Proteins were transferred onto Hybond P PVDF 0.2 membranes (Cytiva) at 10 V for 1 h using a semi-dry blotter (Anatech) with Trans-Blot Turbo transfer buffer (Bio-Rad). The membranes were blocked in Bullet Blocking One (Nacalai Tesque) at room temperature for 10 min and then incubated with mouse monoclonal α-FLAG (1:5,000, Sigma-Aldrich) or rabbit polyclonal α-GroEL (1:10,000, Sigma-Aldrich) antibodies diluted in Bullet Blocking One for 1 h at room temperature or overnight at 4°C. After washing three times for 15 min each with 1×TBST buffer at room temperature, the membranes were incubated with secondary HRP-linked α-mouse or α-rabbit antibodies (1:10,000, Cell Signaling Technology) in Bullet Blocking One for 1 h at room temperature. The membranes were then washed again in three 15-min steps with 1×TBST buffer. Chemiluminescent signals were developed using Amersham ECL Prime reagents (Cytiva), visualized on LAS4000 (GE Healthcare), and quantified using ImageJ software.

### Arginine decarboxylase activity assay

*Salmonella* cells were cultured anaerobically in LB (pH 7.0) until the late exponential phase (OD_660_ = 0.5). The culture pH was then shifted to 5.5 by adding 100 mM MES (pH 5.4). Cells were further cultured for 30 min, harvested by centrifugation (10 min at 3,200 *g*, 20°C) and washed with 0.85% NaCl to normalize cell concentrations. Aliquots of each cell suspension (1.25, 2.5, 5.0, 10, or 20 × 10^7^ cells) were transferred to 200 μl of the arginine decarboxylase assay reagent (1 g L-arginine, 0.05 g bromocresol green, 90 g NaCl, and 3 ml Triton X-100 per liter of distilled water, with the pH adjusted to 3.4 with HCl, (11)). The reaction mixtures were incubated at 37°C for 90 min and qualitatively assessed for decarboxylase activity based on a color change from yellow to blue.

### Growth assay

*Salmonella* cells grown aerobically in LB (pH 7.0) for 21 h were diluted 1:200 in 200 μl of pH 5.5 LB supplemented with 100 mM MES (pH 5.4) in the absence or presence of 1% DMSO, 0.4% glucose, or both in 96-well flat-bottom clear microtiter plates (Iwaki) in a Coy anaerobic chamber (COY). The plates were sealed with SealPlate (Excel Scientific) and incubated at 37℃ with reciprocal shaking at 1,096 rpm and a 1-mm amplitude in a Synergy H1 microplate reader (BioTek). Growth was continuously monitored by measuring OD_600_ every 10 min for 24 h.

### Transcriptional reporter assay

*Salmonella* strains carrying either plasmid pYC964 or its vector control pJV300 were cultured aerobically in LB medium adjusted to pH 7.0, 6.0, or 4.0 with HCl. Cultures were grown to exponential phase, and fluorescence was measured using an Agilent BioTek Synergy H1 plate reader (excitation: 587 nm; emission: 610 nm). RFU was normalized to the corresponding OD_600_ value.

### Single-cell fluorescence imaging and analysis

5 × 10^4^ mouse leukemic monocyte/macrophage cells (RAW264.7; ATCC TIB-71) were seeded in 500 μl of RPMI1640 medium (Thermo Fisher Scientific) supplemented with 10% fetal calf serum (FCS; Biochrom) in a 24-well plate (Corning) and incubated at 37°C in a humidified atmosphere containing 5% CO_2_ for 2 days. A *Salmonella* GFP-reporter strain (*hisG*^+^ *putAP*::P_Llac-O-_*tnaC*_eco_5’UTR-sfGFP-Cm^R^) carrying either pYC964 or its control vector pJV300 was grown overnight in LB medium. The cultures were diluted 1:100 into fresh LB and grown to OD_600_ = 2.0. Bacteria were harvested, washed twice with PBS, resuspended in RPMI medium, and added to the cells at a multiplicity of infection (MOI) of 10. Plates were centrifuged at 250 *g* for 10 min and incubated at 37°C for 30 min. To eliminate extracellular bacteria, the medium was replaced with RPMI containing 50 µg/ml gentamicin for 30 min, followed by RPMI containing 20 µg/ml gentamicin for the remainder of the experiment. At 0, 1, 3, 5, or 20 h.p.i., cells were washed with PBS and fixed with 4% paraformaldehyde for 15 min at room temperature. Nuclei were stained with Hoechst 33258 (Thermo Fisher Scientific). Images were acquired using an Olympus SpinSR10 Ixplore spinning disk confocal microscope equipped with a UPlanApo 60×/1.5 NA oil immersion objective. Image analysis was performed using ImageJ (NIH).

### Macrophage infection assay

2 × 10^5^ RAW264.7 cells were seeded in 2 ml of RPMI1640 medium (Thermo Fisher Scientific) supplemented with 10% FCS (Biochrom), 2 mM L-glutamine (Gibco), and 1 mM sodium pyruvate (Gibco) in a 6-well plate (OmniAb) and incubated at 37°C in a humidified atmosphere containing 5% CO_2_ for 2 days.

*Salmonella* cells were grown aerobically in LB (pH 7.0) for 16 h and harvested by centrifugation (2 min at 10,000 *g*, room temperature). The pellet corresponding to 2 OD_660_ was resuspended in 2 ml of RPMI supplemented with 10% mouse serum (Sigma-Aldrich). For competition assays between two strains, equal volumes of the two suspensions were mixed at a 1:1 ratio. The final mixture was incubated at room temperature for 20 min prior to infection. To check CFU, the mixture was serially diluted in PBS and plated onto LB agar plates. The plates were incubated at 37°C overnight.

RAW264.7 cells were infected by adding 100 μl of the bacterial suspension to each well at a MOI of 25. Following bacterial addition, the plates were centrifuged at 250 *g* for 10 min at room temperature and then incubated at 37°C in a humidified atmosphere containing 5% CO₂ for 30 min. The medium was then replaced with RPMI containing 50 μg/ml gentamicin to eliminate extracellular bacteria, followed by a 30-min incubation. Subsequently, the medium was replaced again with fresh RPMI containing 10 μg/ml gentamicin, and the infected cells were incubated under the same conditions. Time point 0 was defined as the time of the first gentamicin addition.

At 2 or 20 h.p.i., the supernatant was collected, and the infected cells were solubilized using PBS containing 0.1% Triton X-100 (Gibco). Both the supernatant and lysate samples were serially diluted in PBS and plated onto LB agar plates. The plates were incubated at 37°C overnight. Fluorescence signals from sfGFP or mSc within the colonies were detected using a trans illuminator (Nippon Genetics) to determine the CFU for each strain.

## ACKNOWLEDGMENTS

We thank all members of the Westermann, Chao, and Miyakoshi labs for their helpful discussions and support throughout the project. We also thank Teppei Morita (Kyorin University) for his valuable comments.

---

**Figure S1.**
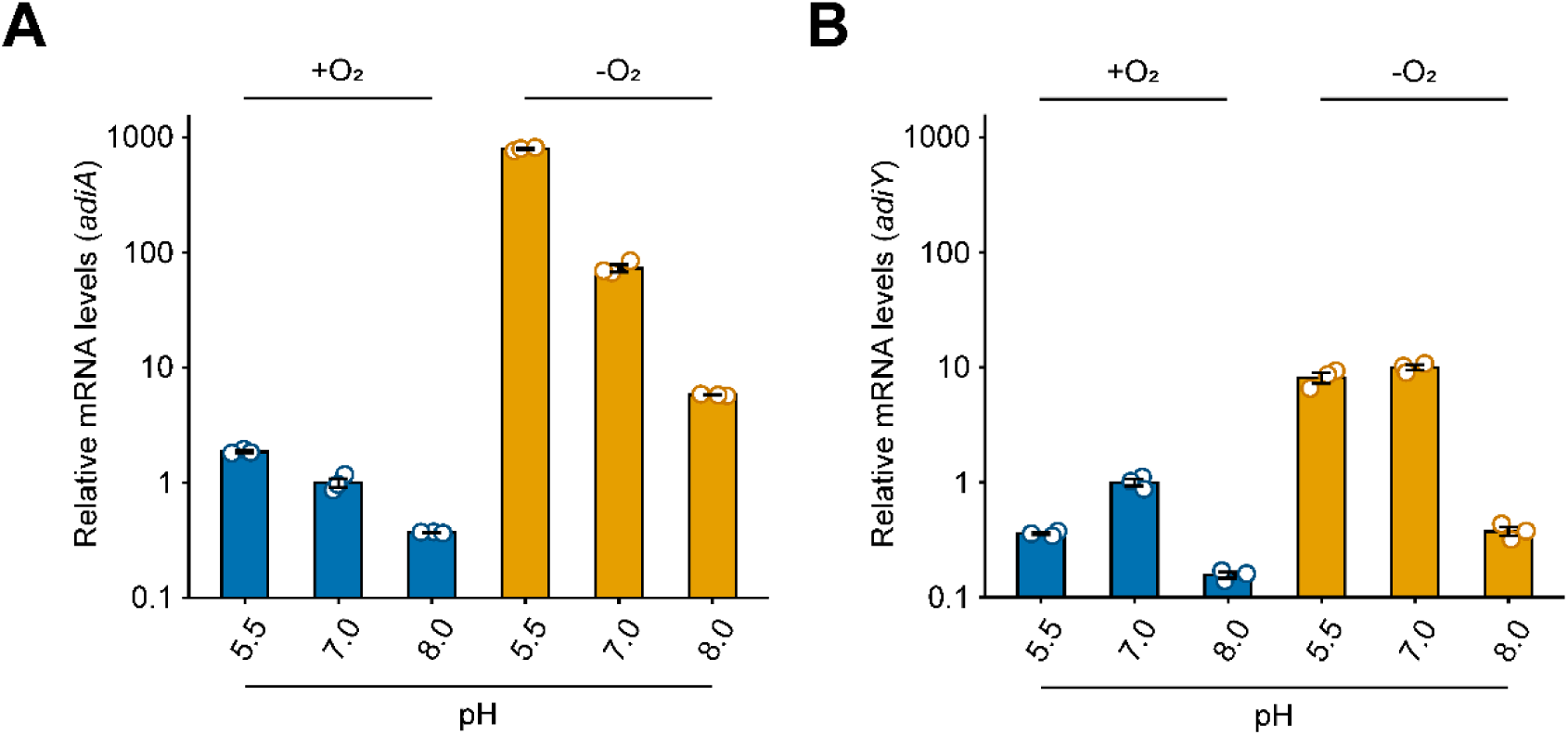
Expression profiles of *adiA* (A) and *adiY* (B). Wild-type *Salmonella* was cultured at pH 7.0 under aerobic or anaerobic conditions until the late exponential phase. The culture pH was shifted to acidic (pH 5.5), alkaline (pH 8.0), or maintained at neutral (pH 7.0). Total RNA extracted 30 min after the pH shift was analyzed by qRT-PCR. Relative mRNA levels of *adiA* and *adiY* were normalized to those of *rpoB*, with the levels at pH 7.0 under aerobic conditions set to 1. Data are presented as mean ± standard error from three independent experiments (n = 3).

**Figure S2.**
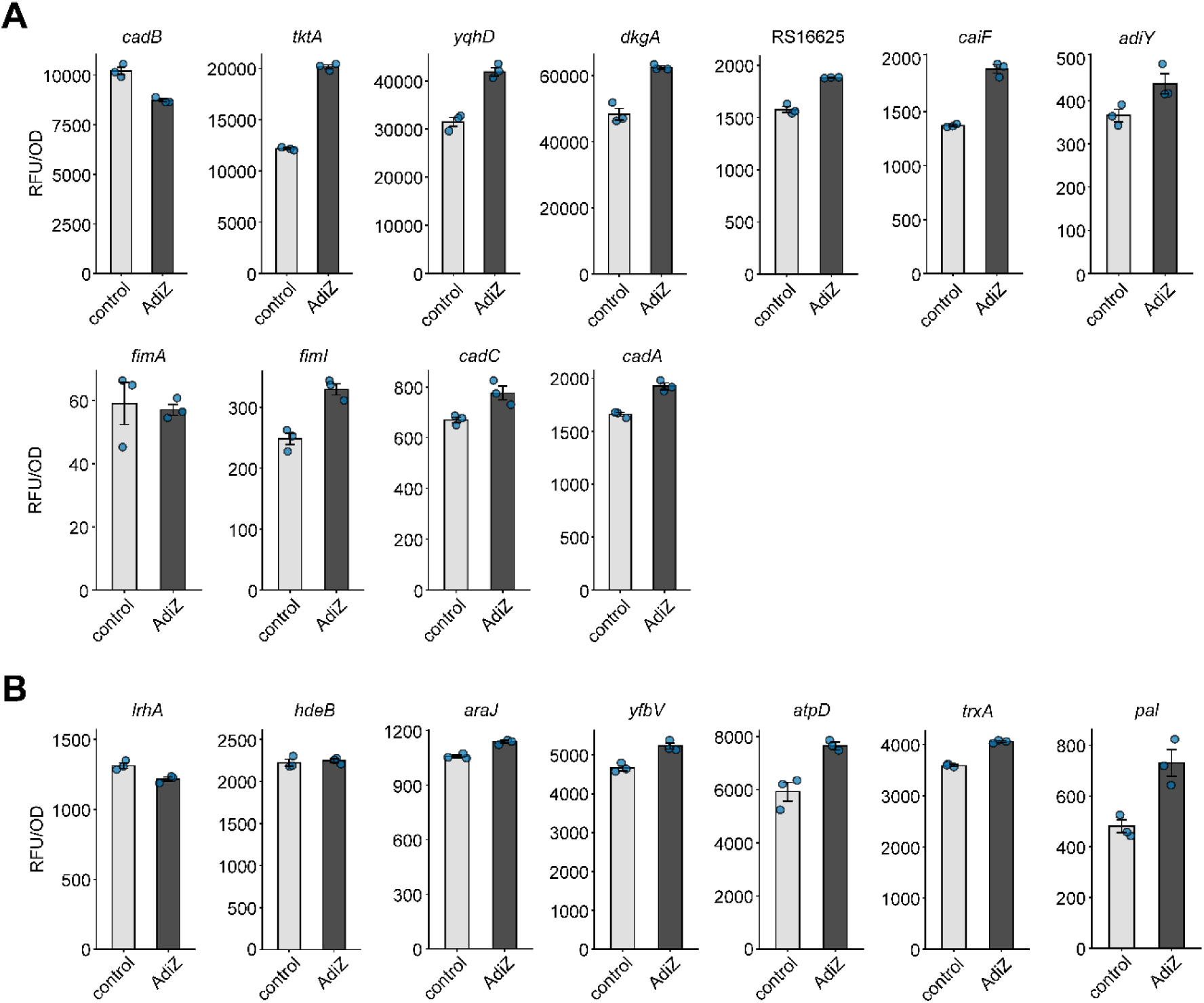
sfGFP-reporter assay. AdiZ target candidates identified by RNA-seq following AdiZ pulse-expression (A) and iRIL-seq (B) were analyzed by sfGFP-reporter assay. *E*. *coli* Δ*adiZ* strains harboring a combination of the sfGFP-reporter plasmid and either pJV300 (vector control) or pP_L_-AdiZ were cultured aerobically to the exponential phase. RFU was calculated by subtracting the autofluorescence of the same strains without sfGFP-reporter plasmids and normalized to OD_600_ value. Data are presented as mean ± standard error from biological triplicates (n = 3).

**Figure S3.**
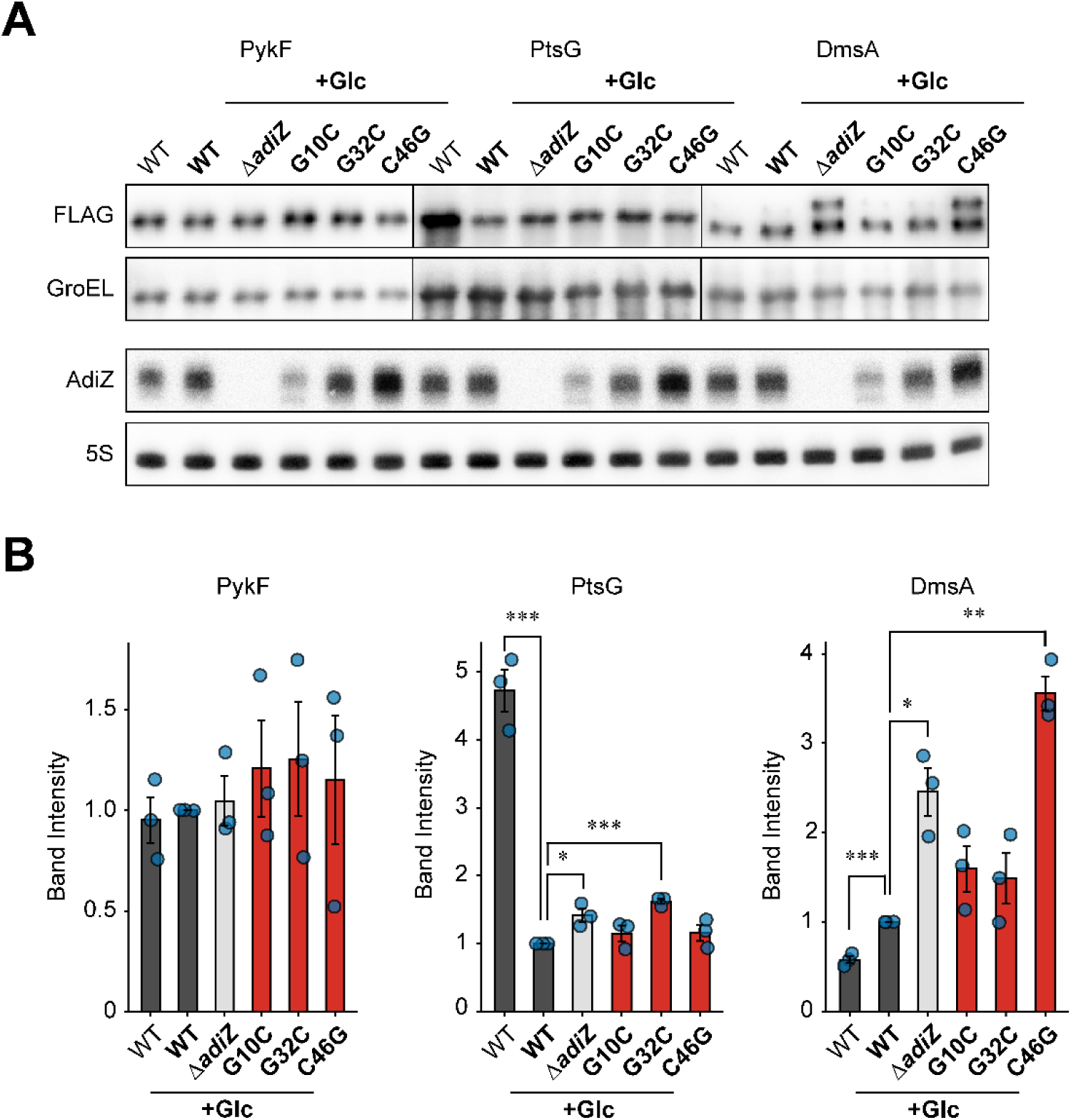
Western blot analysis of endogenous protein levels of AdiZ targets in the presence of glucose. (A) Wild-type *Salmonella*, the Δ*adiZ* mutant, and the chromosomal point mutants (*adiZ*-G10C, G32C, or C46G) with C-terminal 3x-FLAG fusions to PykF, PtsG, or DmsA were cultured anaerobically in LB (pH 5.5) supplemented with 0.4% glucose until the late exponential phase. RNA extracted under the same conditions was analyzed by northern blot. AdiZ was detected with MMO-2143 and 5S rRNA detected with MMO-1056 served as a loading control. A representative image from three independent experiments is shown. (B) Band intensities of PykF, PtsG, and DmsA normalized to GroEL. Data are presented as mean ± standard error (n = 3). Statistical significance was assessed using one-way ANOVA followed by two-tailed Student’s t-test with Bonferroni correction (*p < 0.05, **p < 0.01, ***p < 0.001).

**Figure S4.**
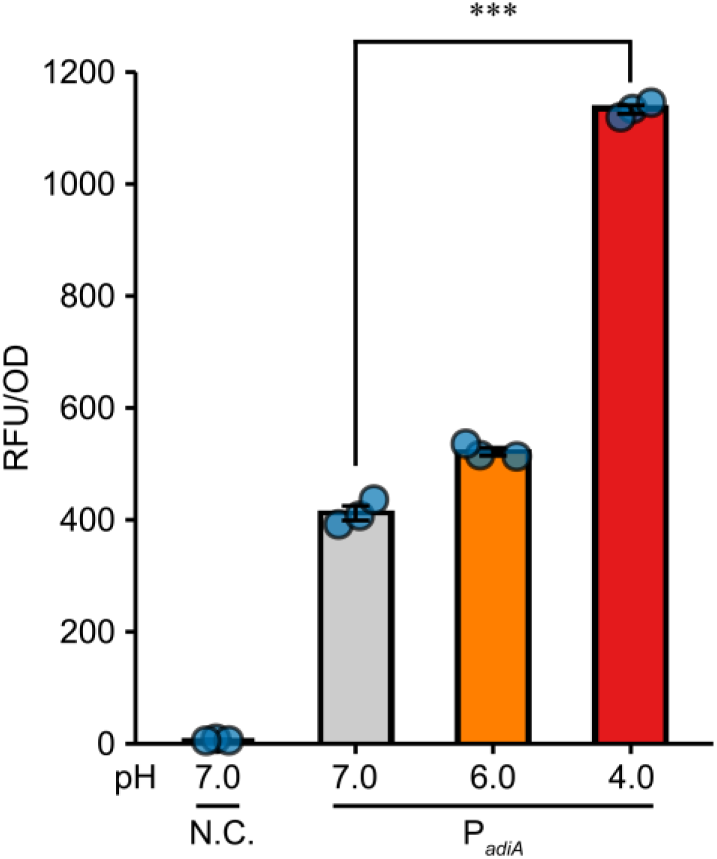
Transcriptional reporter of *adiA* is activated under acidic conditions. A *Salmonella* strain harboring the P*_adiA_*-mCherry transcriptional reporter plasmid (pYC964) was cultured aerobically in LB at pH 7.0, 6.0, or 4.0 until the exponential phase. A strain harboring the vector control (pJV300) cultured at pH 7.0 served as a negative control. RFU was calculated by subtracting the autofluorescence of the same strains without the reporter plasmids and normalized to OD_600_ value. Data are presented as mean ± standard error from biological triplicates (n = 3). Statistical significance was analyzed using one-way ANOVA with Dunnett’s test (***p < 0.001).

**Figure S5.**
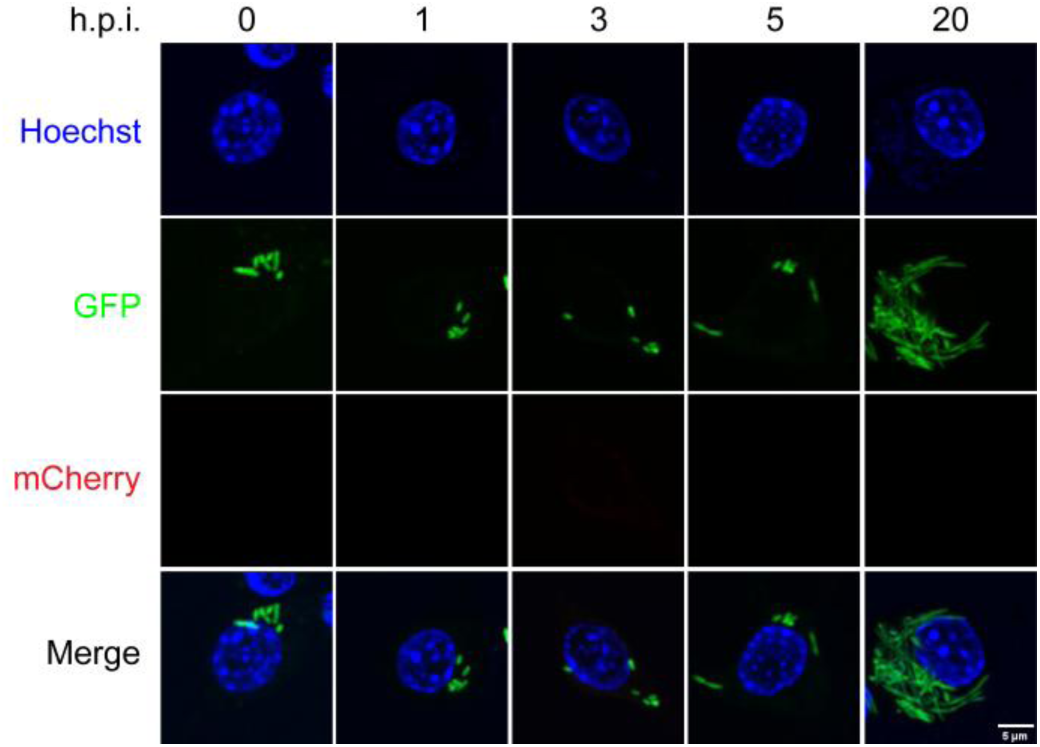
Fluorescence microscopic observation of the *Salmonella* control strain for the transcriptional reporter. An sfGFP-expressing *Salmonella* strain harboring the pJV300 empty vector control for the transcriptional reporter was added to cultures of mouse macrophage-like RAW264.7 cells. At the indicated time points, cells were fixed, stained with Hoechst, and analyzed by fluorescence microscopy. Representative images are shown from two independent biological replicates.

**Figure S6.**
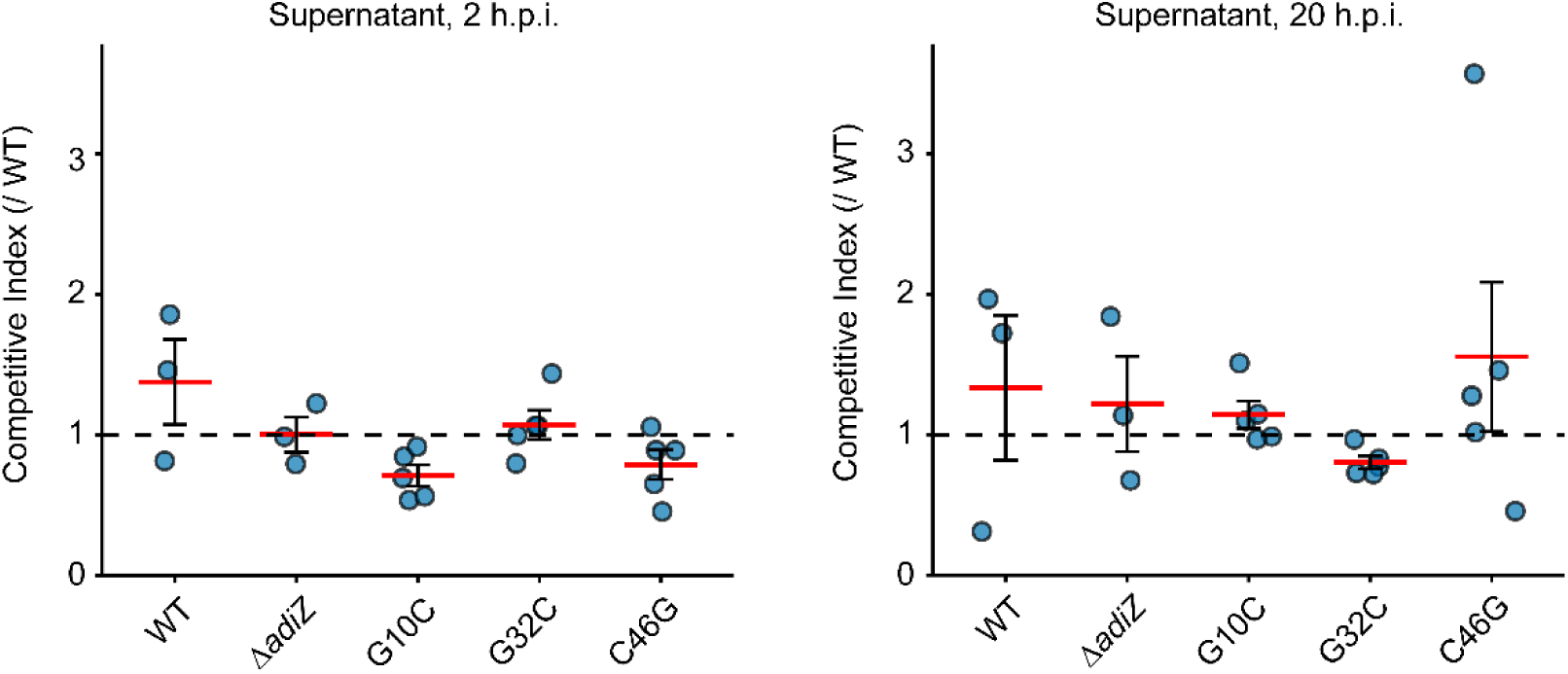
CI values in the supernatant of macrophage cell culture. The dashed line represents CI = 1. Red bars represent the mean CI with SEM (n = 3 or 5). According to a two-tailed Student’s t-test, none of the differences were statistically significant.

**Figure S7.**
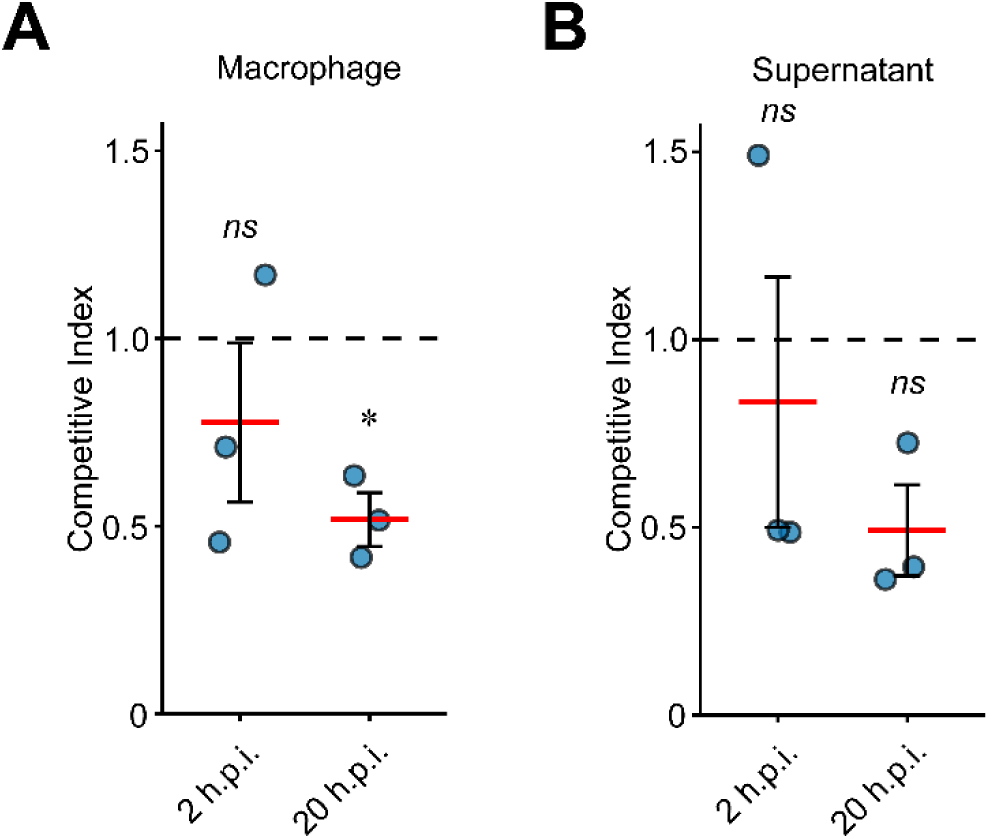
Competitive infection assay between mSc *adiZ*-WT and sfGFP Δ*adiZ*. CI values within macrophage cells (A) and in the culture supernatant (B). The dashed line represents CI = 1. Red bars represent the mean CI with SEM (n = 3). Statistical significance was determined using the two-tailed Student’s t-test. p < 0.05, “*ns*” indicates no significant difference.

**Figure S8.**
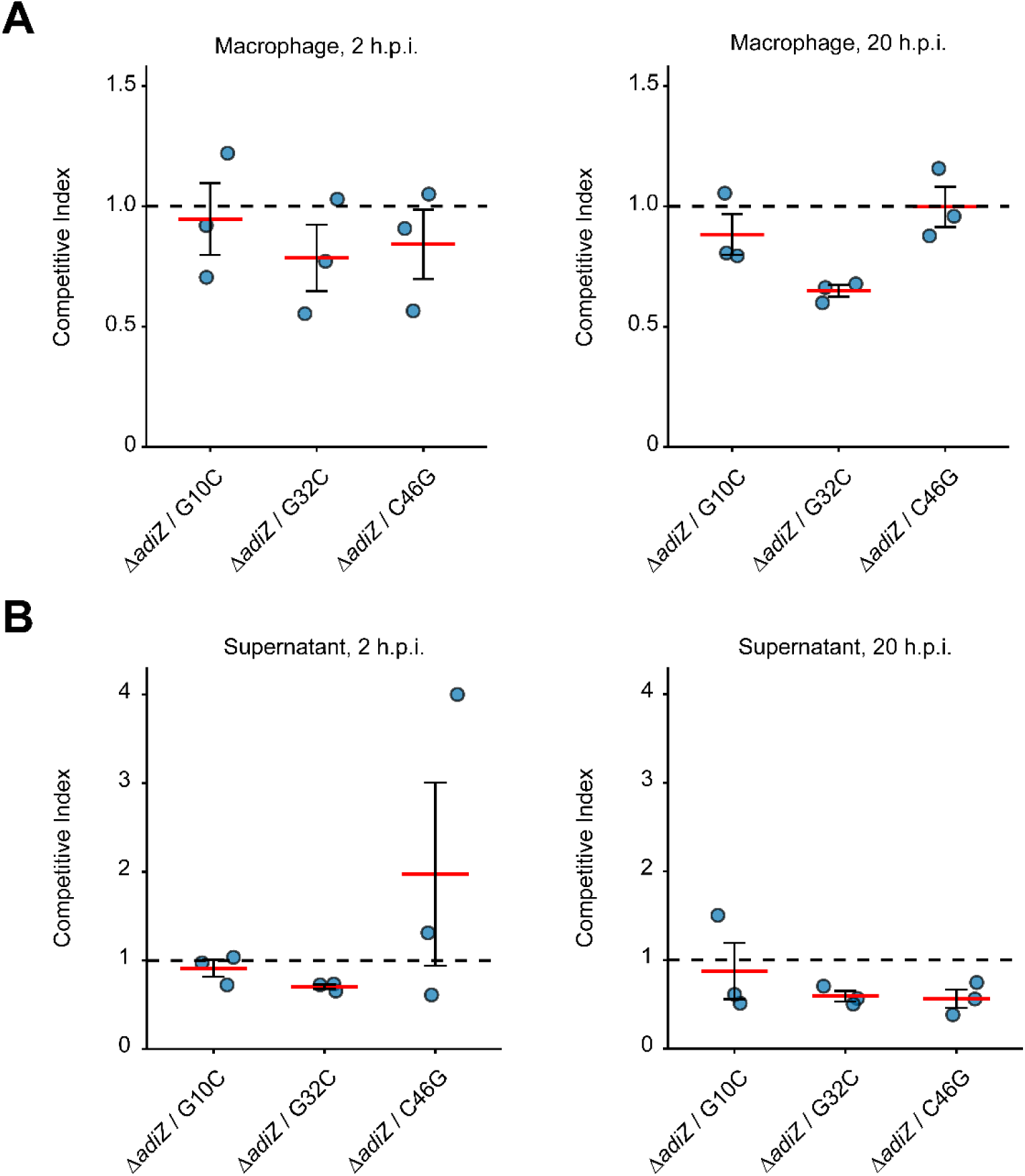
Competitive infection assay between sfGFP Δ*adiZ* and mSc *adiZ* point mutants. CI values within macrophage cells (A) and in the culture supernatant (B). The dashed line represents CI = 1. Red bars represent the mean CI with SEM (n = 3). According to a two-tailed Student’s t-test, none of the differences were statistically significant.

**Figure S9.**
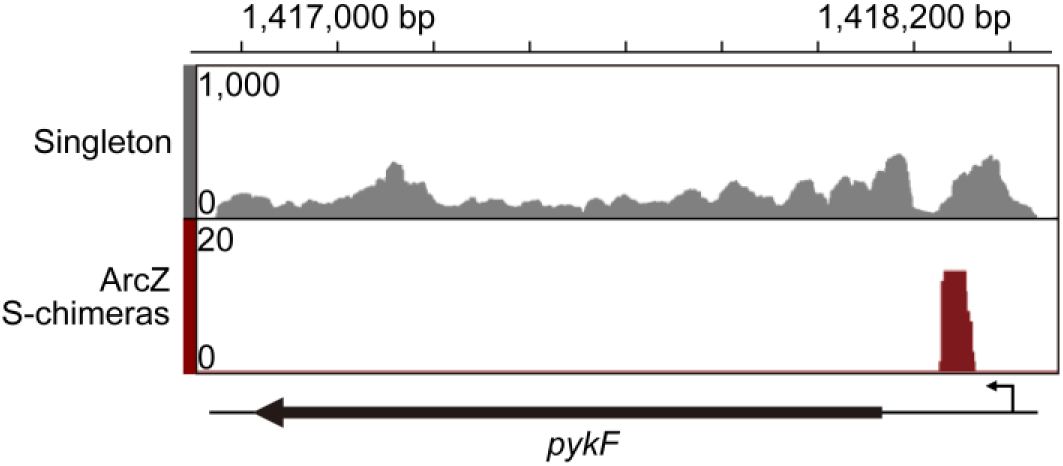
Genome browser view of iRIL-seq reads showing the interaction between ArcZ and pykF. A representative screenshot displaying singleton and chimeric reads from iRIL-seq, with the ArcZ sRNA mapped to the 5′UTR of *pykF*.

**Figure S10.**
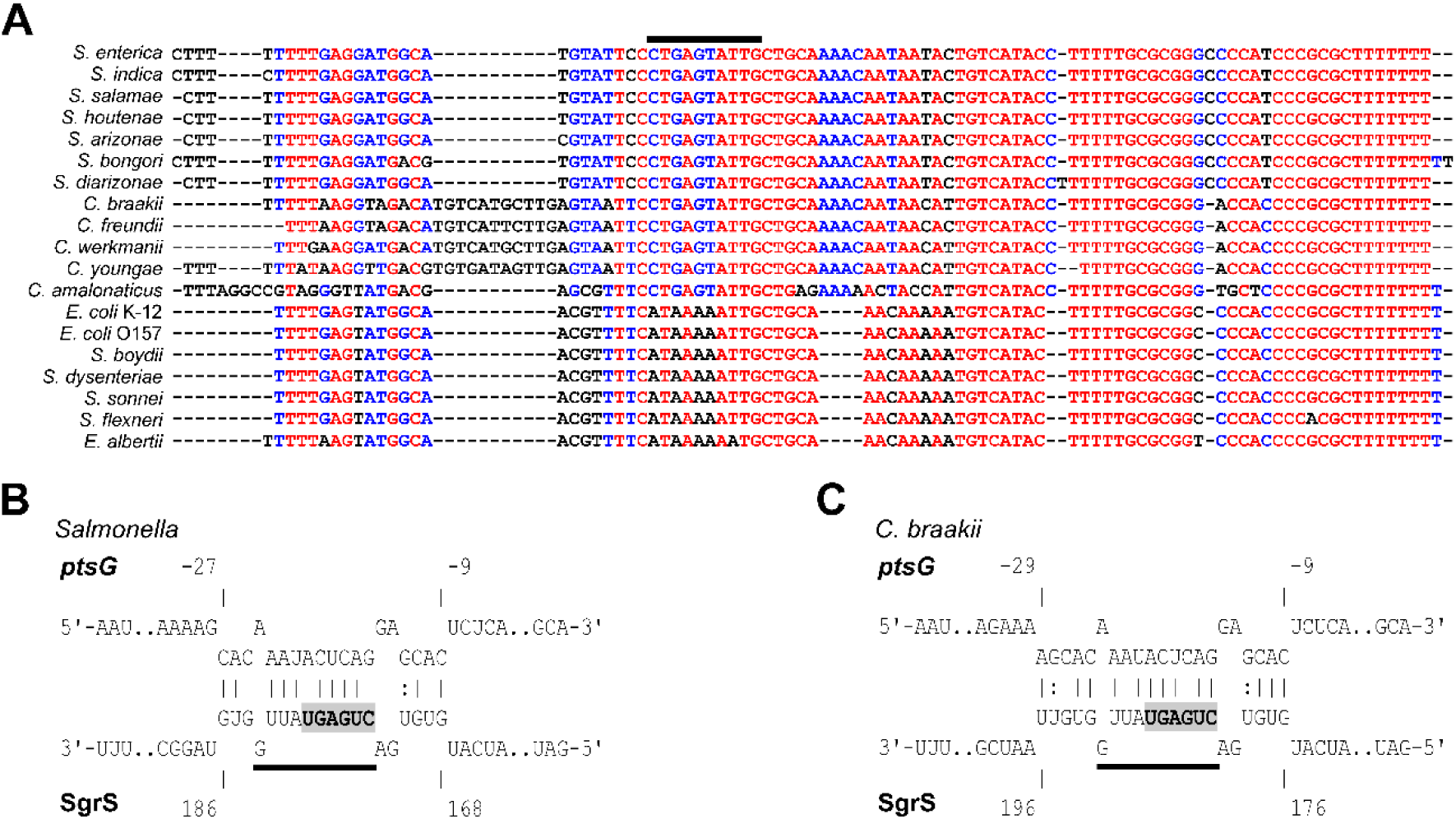
AdiZ and SgrS share critical sequence motifs for base-pairing with *ptsG* mRNA in Salmonella *and* Citrobacter. (A) Multiple sequence alignment of *adiZ,* as shown in Figure 1C. (B, C) Base-pairing interactions between SgrS and *ptsG* mRNA in *Salmonella* (B) and *C. braakii* (C) predicted by the IntaRNA program. Numbers above and below the nucleotide sequences indicate the position relative to the start codon of *ptsG* and the transcription start site of SgrS, respectively. Shared nucleotide sequences between AdiZ and SgrS required for base-pairing with *ptsG* mRNA are indicated by a solid line. The six nucleotides forming the critical core (58) are highlighted in bold.

## REFERENCES

1. Majowicz SE, Musto J, Scallan E, Angulo FJ, Kirk M, O’Brien SJ, Jones TF, Fazil A, Hoekstra RM. 2010. The global burden of nontyphoidal salmonella gastroenteritis. Clinical Infectious Diseases 50:882–889.

2. Stanaway JD, Reiner RC, Blacker BF, Goldberg EM, Khalil IA, Troeger CE, Andrews JR, Bhutta ZA, Crump JA, Im J, Marks F, Mintz E, Park SE, Zaidi AKM, Abebe Z, Abejie AN, Adedeji IA, Ali BA, Amare AT, Atalay HT, Avokpaho EFGA, Bacha U, Barac A, Bedi N, Berhane A, Browne AJ, Chirinos JL, Chitheer A, Dolecek C, El Sayed Zaki M, Eshrati B, Foreman KJ, Gemechu A, Gupta R, Hailu GB, Henok A, Hibstu DT, Hoang CL, Ilesanmi OS, Iyer VJ, Kahsay A, Kasaeian A, Kassa TD, Khan EA, Khang YH, Magdy Abd El Razek H, Melku M, Mengistu DT, Mohammad KA, Mohammed S, Mokdad AH, Nachega JB, Naheed A, Nguyen CT, Nguyen HLT, Nguyen LH, Nguyen NB, Nguyen TH, Nirayo YL, Pangestu T, Patton GC, Qorbani M, Rai RK, Rana SM, Ranabhat CL, Roba KT, Roberts NLS, Rubino S, Safiri S, Sartorius B, Sawhney M, Shiferaw MS, Smith DL, Sykes BL, Tran BX, Tran TT, Ukwaja KN, Vu GT, Vu LG, Weldegebreal F, Yenit MK, Murray CJL, Hay SI. 2019. The global burden of typhoid and paratyphoid fevers: a systematic analysis for the Global Burden of Disease Study 2017. Lancet Infect Dis 19:369–381.

3. Ao TT, Feasey NA, Gordon MA, Keddy KH, Angulo FJ, Crump JA. 2015. Global burden of invasive nontyphoidal salmonella disease, 2010. Emerg Infect Dis 21:941–949.

4. Smith JL. 2003. The role of gastric acid in preventing foodborne disease and how bacteria overcome acid conditions. J Food Prot 66:1292–1303.

5. Alpuche Aranda CM, Swanson JA, Loomis WP, Miller SI. 1992. Salmonella typhimurium activates virulence gene transcription within acidified macrophage phagosomes. Proc Natl Acad Sci U S A 89:10079–10083.

6. Gong S, Richard H, Foster JW. 2003. YjdE (AdiC) is the arginine:Agmatine antiporter essential for arginine-dependent acid resistance in Escherichia coli. J Bacteriol 185:4402–4409.

7. Foster JW. 2004. Escherichia coli acid resistance: Tales of an amateur acidophile. Nat Rev Microbiol 2:898–907.

8. Radlinski LC, Rogers AWL, Bechtold L, Masson HLP, Nguyen H, Larabi AB, Tiffany CR, Carvalho TP de, Tsolis RM, Bäumler AJ. 2024. Salmonella virulence factors induce amino acid malabsorption in the ileum to promote ecosystem invasion of the large intestine. Proc Natl Acad Sci U S A 121.

9. Viala JPM, Méresse S, Pocachard B, Guilhon AA, Aussel L, Barras F. 2011. Sensing and adaptation to low pH mediated by inducible amino acid decarboxylases in Salmonella. PLoS One 6:e22397.

10. Kieboom J, Abee T. 2006. Arginine-dependent acid resistance in Salmonella enterica serovar typhimurium. J Bacteriol 188:5650–5653.

11. Brenneman KE, Willingham C, Kong W, Curtiss R, Roland KL. 2013. Low-ph rescue of acid-sensitive salmonella enterica serovar typhi strains by a rhamnose-regulated arginine decarboxylase system. J Bacteriol 195:3062–3072.

12. Ishikawa H, Otaka H, Maki K, Morita T, Aiba H. 2012. The functional Hfq-binding module of bacterial sRNAs consists of a double or single hairpin preceded by a U-rich sequence and followed by a 3′ poly(U) tail. RNA 18:1062–1074.

13. Kröger C, Dillon SC, Cameron ADS, Papenfort K, Sivasankaran SK, Hokamp K, Chao Y, Sittka A, Hébrard M, Händler K, Colgan A, Leekitcharoenphon P, Langridge GC, Lohan AJ, Loftus B, Lucchini S, Ussery DW, Dorman CJ, Thomson NR, Vogel J, Hinton JCD. 2012. The transcriptional landscape and small RNAs of Salmonella enterica serovar Typhimurium. Proc Natl Acad Sci U S A 109:1277–1286.

14. Kröger C, Colgan A, Srikumar S, Händler K, Sivasankaran SK, Hammarlöf DL, Canals R, Grissom JE, Conway T, Hokamp K, Hinton JCD. 2013. An infection-relevant transcriptomic compendium for salmonella enterica serovar typhimurium. Cell Host Microbe 14:683–695.

15. Canals R, Hammarlöf DL, Kröger C, Owen S V., Fong WY, Lacharme-Lora L, Zhu X, Wenner N, Carden SE, Honeycutt J, Monack DM, Kingsley RA, Brownridge P, Chaudhuri RR, Rowe WPM, Predeus A V., Hokamp K, Gordon MA, Hinton JCD. 2019. Adding function to the genome of African Salmonella Typhimurium ST313 strain D23580. PLoS Biol 17:1–32.

16. Storz G, Vogel J, Wassarman KM. 2011. Regulation by Small RNAs in Bacteria: Expanding Frontiers. Mol Cell 43:880–891.

17. Hör J, Matera G, Vogel J, Gottesman S, Storz G. 2020. Trans-Acting Small RNAs and Their Effects on Gene Expression in Escherichia coli and Salmonella enterica. EcoSal Plus 9.

18. Zhang S, Chao Y. 2025. Quality over quantity: Small RNA pauses translation elongation to lift protein activity. Mol Cell. Cell Press 10.1016/j.molcel.2025.04.003.

19. Miyakoshi M, Chao Y, Vogel J. 2015. Regulatory small RNAs from the 3’ regions of bacterial mRNAs. Curr Opin Microbiol. Elsevier Current Trends 10.1016/j.mib.2015.01.013.

20. Ponath F, Hör J, Vogel J. 2022. An overview of gene regulation in bacteria by small RNAs derived from mRNA 3’ ends. FEMS Microbiol Rev 46:1–18.

21. Papenfort K, Storz G. 2024. Insights into bacterial metabolism from small RNAs. Cell Chem Biol. Cell Press 10.1016/j.chembiol.2024.07.002.

22. Chen Z, Yang Y, Chen X, Bei C, Gao Q, Chao Y, Wang C. 2025. An RNase III– processed sRNA coordinates sialic acid metabolism of Salmonella enterica during gut colonization. Proc Natl Acad Sci U S A 122.

23. De Mets F, Van Melderen L, Gottesman S. 2019. Regulation of acetate metabolism and coordination with the TCA cycle via a processed small RNA. Proc Natl Acad Sci U S A 116:1043–1052.

24. Miyakoshi M, Matera G, Maki K, Sone Y, Vogel J. 2019. Functional expansion of a TCA cycle operon mRNA by a 3 end-derived small RNA. Nucleic Acids Res 47:2075–2088.

25. Walling LR, Kouse AB, Shabalina SA, Zhang H, Storz G. 2022. A 3 UTR-derived small RNA connecting nitrogen and carbon metabolism in enteric bacteria. Nucleic Acids Res 50:10093–10109.

26. Miyakoshi M, Morita T, Kobayashi A, Berger A, Takahashi H, Gotoh Y, Hayashi T, Tanaka K. 2022. Glutamine synthetase mRNA releases sRNA from its 3′UTR to regulate carbon/nitrogen metabolic balance in Enterobacteriaceae. Elife 11:1–24.

27. Iosub IA, Marchioretto M, van Nues RW, McKellar S, Viero G, Granneman S. 2021. The mRNA derived MalH sRNA contributes to alternative carbon source utilization by tuning maltoporin expression in E. coli. RNA Biol 18:914–931.

28. Wang C, Chao Y, Matera G, Gao Q, Vogel J. 2020. The conserved 3′ UTR-derived small RNA NarS mediates mRNA crossregulation during nitrate respiration. Nucleic Acids Res 48:2126–2143.

29. Chao Y, Vogel J. 2016. A 3’ UTR-Derived Small RNA Provides the Regulatory Noncoding Arm of the Inner Membrane Stress Response. Mol Cell 61:352–363.

30. Papenfort K, Melamed S. 2023. Small RNAs, Large Networks: Posttranscriptional Regulons in Gram-Negative Bacteria. Annu Rev Microbiol 77:23–43.

31. Liu F, Chen Z, Zhang S, Wu K, Bei C, Wang C, Chao Y. 2023. In vivo RNA interactome profiling reveals 3’UTR-processed small RNA targeting a central regulatory hub. Nat Commun 14.

32. Wu K, Lin X, Lu Y, Dong R, Jiang H, Svensson SL, Zheng J, Shen N, Camilli A, Chao Y. 2024. RNA interactome of hypervirulent Klebsiella pneumoniae reveals a small RNA inhibitor of capsular mucoviscosity and virulence. Nat Commun 15:1–15.

33. Enriqueta Muñoz M, Ponce E. 2003. Pyruvate kinase: Current status of regulatory and functional properties. Comparative Biochemistry and Physiology - B Biochemistry and Molecular Biology 135:197–218.

34. Postma PW, Lengeler JW, Jacobson GR. 1993. Phosphoenolpyruvate : Carbohydrate phosphotransferase systems of bacteria. Microbiol Rev 57:543–594.

35. Kim JS, Liu L, Kant S, Orlicky DJ, Uppalapati S, Margolis A, Davenport BJ, Morrison TE, Matsuda J, McClelland M, Jones-Carson J, Vazquez-Torres A. 2024. Anaerobic respiration of host-derived methionine sulfoxide protects intracellular Salmonella from the phagocyte NADPH oxidase. Cell Host Microbe 32:411–424.e10.

36. Brameyer S, Schumacher K, Kuppermann S, Jung K. 2022. Division of labor and collective functionality in Escherichia coli under acid stress. Commun Biol 5:1–14.

37. Zuker M. 2003. Mfold web server for nucleic acid folding and hybridization prediction. Nucleic Acids Res 31:3406–3415.

38. Chao Y, Li L, Girodat D, Förstner KU, Said N, Corcoran C, Śmiga M, Papenfort K, Reinhardt R, Wieden HJ, Luisi BF, Vogel J. 2017. In Vivo Cleavage Map Illuminates the Central Role of RNase E in Coding and Non-coding RNA Pathways. Mol Cell 65:39–51.

39. Holmqvist E, Wright PR, Li L, Bischler T, Barquist L, Reinhardt R, Backofen R, Vogel J. 2016. Global RNA recognition patterns of post-transcriptional regulators Hfq and CsrA revealed by UV crosslinking in vivo . EMBO J 35:991–1011.

40. Lee Catherine A, Jones BD, Falkow S. 1992. Identification of a Salmonella typhimurium invasion locus by selection for hyperinvasive mutants. Proc Natl Acad Sci U S A 89:1847–1851.

41. Mann M, Wright PR, Backofen R. 2017. IntaRNA 2.0: Enhanced and customizable prediction of RNA-RNA interactions. Nucleic Acids Res 45:W435–W439.

42. Corcoran CP, Podkaminski D, Papenfort K, Urban JH, Hinton JCD, Vogel J. 2012. Superfolder GFP reporters validate diverse new mRNA targets of the classic porin regulator, MicF RNA. Mol Microbiol 84:428–445.

43. Sambasivarao D, Turner RJ, Simala-Grant JL, Shaw G, Hu J, Weiner JH. 2000. Multiple roles for the twin arginine leader sequence of dimethyl sulfoxide reductase of Escherichia coli. Journal of Biological Chemistry 275:22526–22531.

44. Bilous PT, Cole ST, Anderson WF, Weiner JH. 1988. Nucleotide sequence of the dmsABC operon encoding the anaerobic dimethylsulphoxide reductase of Escherichia coli. Mol Microbiol 2:785–795.

45. Kimata K, Takahashi H, Inada T, Postma P, Aiba H. 1997. cAMP receptor protein-cAMP plays a crucial role in glucose-lactose diauxie by activating the major glucose transporter gene in Escherichia coli. Proc Natl Acad Sci U S A 94:12914–12919.

46. Vanderpool CK, Gottesman S. 2004. Involvement of a novel transcriptional activator and small RNA in post-transcriptional regulation of the glucose phosphoenolpyruvate phosphotransferase system. Mol Microbiol 54:1076–1089.

47. Morita T, Mochizuki Y, Aiba H. 2006. Translational repression is sufficient for gene silencing by bacterial small noncoding RNAs in the absence of mRNA destruction. Proc Natl Acad Sci U S A 103:4858–4863.

48. Chakraborty S, Liu L, Fitzsimmons L, Porwollik S, Kim JS, Desai P, McClelland M, Vazquez-Torres A. 2020. Glycolytic reprograming in Salmonella counters NOX2-mediated dissipation of ΔpH. Nat Commun 11:1–11.

49. Nguyen TH, Wang BX, Diaz OR, Rajendram M, Mckenna JA, Butler DSC, Hokamp K, Hinton JCD, Monack DM, Kerwyn &, Huang C. 2025. Profiling Salmonella transcriptional dynamics during macrophage infection using a comprehensive reporter library. Nature Microbiology 2025 10:4 10:1006–1023.

50. Melamed S, Peer A, Faigenbaum-Romm R, Gatt YE, Reiss N, Bar A, Altuvia Y, Argaman L, Margalit H. 2016. Global Mapping of Small RNA-Target Interactions in Bacteria. Mol Cell 63:884–897.

51. Melamed S, Adams PP, Zhang A, Zhang H, Storz G. 2020. RNA-RNA Interactomes of ProQ and Hfq Reveal Overlapping and Competing Roles. Mol Cell 77:411–425.e7.

52. Iosub IA, van Nues RW, McKellar SW, Nieken KJ, Marchioretto M, Sy B, Tree JJ, Viero G, Granneman S. 2020. Hfq CLASH uncovers sRNA-target interaction networks linked to nutrient availability adaptation. Elife 9:1–33.

53. Mizrahi SP, Elbaz N, Argaman L, Altuvia Y, Katsowich N, Socol Y, Bar A, Rosenshine I, Margalit H. 2021. The impact of Hfq-mediated sRNA-mRNA interactome on the virulence of enteropathogenic Escherichia coli. Sci Adv 7:1–14.

54. Matera G, Altuvia Y, Gerovac M, El Mouali Y, Margalit H, Vogel J. 2022. Global RNA interactome of Salmonella discovers a 5′ UTR sponge for the MicF small RNA that connects membrane permeability to transport capacity. Mol Cell 82:629–644.e4.

55. McQuail J, Matera G, Gräfenhan T, Bischler T, Haberkant P, Stein F, Vogel J, Wigneshweraraj S. 2024. Global Hfq-mediated RNA interactome of nitrogen starved Escherichia coli uncovers a conserved post-transcriptional regulatory axis required for optimal growth recovery. Nucleic Acids Res 52:2323–2339.

56. Westermann AJ. 2018. Regulatory RNAs in Virulence and Host-Microbe Interactions. Microbiol Spectr 6.

57. Srikumar S, Kröger C, Hébrard M, Colgan A, Owen S V., Sivasankaran SK, Cameron ADS, Hokamp K, Hinton JCD. 2015. RNA-seq Brings New Insights to the Intra-Macrophage Transcriptome of Salmonella Typhimurium. PLoS Pathog 11:e1005262.

58. Kawamoto H, Koide Y, Morita T, Aiba H. 2006. Base-pairing requirement for RNA silencing by a bacterial small RNA and acceleration of duplex formation by Hfq. Mol Microbiol 61:1013–1022.

59. Papenfort K, Vogel J. 2014. Small RNA functions in carbon metabolism and virulence of enteric pathogens. Front Cell Infect Microbiol 4:1–12.

60. Westermann AJ, Förstner KU, Amman F, Barquist L, Chao Y, Schulte LN, Müller L, Reinhardt R, Stadler PF, Vogel J. 2016. Dual RNA-seq unveils noncoding RNA functions in host-pathogen interactions. Nature 529:496–501.

61. Kim K, Palmer AD, Vanderpool CK, Slauch JM. 2019. The small RNA pint contributes to phop-mediated regulation of the salmonella pathogenicity island 1 type III secretion system in salmonella enterica serovar typhimurium. J Bacteriol 201.

62. Ryan D, Mukherjee M, Nayak R, Dutta R, Suar M. 2018. Biological and regulatory roles of acid-induced small RNA RyeC in Salmonella Typhimurium. Biochimie 150:48–56.

63. Papenfort K, Podkaminski D, Hinton JCD, Vogel J. 2012. The ancestral SgrS RNA discriminates horizontally acquired Salmonella mRNAs through a single G-U wobble pair. Proc Natl Acad Sci U S A 109.

64. Bobrovskyy M, Vanderpool CK. 2016. Diverse mechanisms of post-transcriptional repression by the small RNA regulator of glucose-phosphate stress. Mol Microbiol 99:254–273.

65. Kenney LJ. 2019. The role of acid stress in Salmonella pathogenesis. Curr Opin Microbiol 47:45–51.

66. Liew ATF, Foo YH, Gao Y, Zangoui P, Singh MK, Gulvady R, Kenney LJ. 2019. Single cell, super-resolution imaging reveals an acid pH-dependent conformational switch in SsrB regulates SPI-2. Elife 8:1–26.

67. Shetty D, Kenney LJ. 2023. A pH-sensitive switch activates virulence in Salmonella. Elife 12:1–20.

68. Chakraborty S, Mizusaki H, Kenney LJ. 2015. A FRET-Based DNA Biosensor Tracks OmpR-Dependent Acidification of Salmonella during Macrophage Infection. PLoS Biol 13:1–32.

69. Steeb B, Claudi B, Burton NA, Tienz P, Schmidt A, Farhan H, Mazé A, Bumann D. 2013. Parallel Exploitation of Diverse Host Nutrients Enhances Salmonella Virulence. PLoS Pathog 9.

70. Margolis A, Liu L, Porwollik S, Till JKA, Chu W, McClelland M, Vázquez-Torres A. 2023. Arginine Metabolism Powers Salmonella Resistance to Oxidative Stress. Infect Immun 91.

71. Das P, Lahiri A, Lahiri A, Sen M, Iyer N, Kapoor N, Balaji KN, Chakravortty D. 2010. Cationic Amino Acid Transporters and Salmonella Typhimurium ArgT Collectively Regulate Arginine Availability towards Intracellular Salmonella Growth. PLoS One 5.

72. Choi Y, Choi J, Groisman EA, Kang DH, Shin D, Ryu S. 2012. Expression of stm4467-encoded arginine deiminase controlled by the stm4463 regulator contributes to salmonella enterica serovar typhimurium virulence. Infect Immun 80:4291–4297.

73. Miki T, Uemura T, Kinoshita M, Ami Y, Ito M, Okada N, Furuchi T, Kurihara S, Haneda T, Minamino T, Kim YG. 2024. Salmonella Typhimurium exploits host polyamines for assembly of the type 3 secretion machinery. PLoS Biol 22:e3002731.

74. Doherty GP, Bailey K, Lewis PJ. 2010. Stage-specific fluorescence intensity of GFP and mCherry during sporulation in Bacillus Subtilis. BMC Res Notes 3:1–8.

75. Eriksson S, Lucchini S, Thompson A, Rhen M, Hinton JCD. 2003. Unravelling the biology of macrophage infection by gene expression profiling of intracellular Salmonella enterica. Mol Microbiol 47:103–118.

76. Chaudhuri RR, Morgan E, Peters SE, Pleasance SJ, Hudson DL, Davies HM, Wang J, van Diemen PM, Buckley AM, Bowen AJ, Pullinger GD, Turner DJ, Langridge GC, Turner AK, Parkhill J, Charles IG, Maskell DJ, Stevens MP. 2013. Comprehensive Assignment of Roles for Salmonella Typhimurium Genes in Intestinal Colonization of Food-Producing Animals. PLoS Genet 9.

77. Papenfort K, Pfeiffer V, Mika F, Lucchini S, Hinton JCD, Vogel J. 2006. σE-dependent small RNAs of Salmonella respond to membrane stress by accelerating global omp mRNA decay. Mol Microbiol 62:1674–1688.

78. Sittka A, Pfeiffer V, Tedin K, Vogel J. 2007. The RNA chaperone Hfq is essential for the virulence of Salmonella typhimurium. Mol Microbiol 63:193–217.

79. Urban JH, Vogel J. 2007. Translational control and target recognition by Escherichia coli small RNAs in vivo. Nucleic Acids Res 35:1018–1037.

80. Bindels DS, Haarbosch L, Van Weeren L, Postma M, Wiese KE, Mastop M, Aumonier S, Gotthard G, Royant A, Hink MA, Gadella TWJ. 2016. mScarlet: A bright monomeric red fluorescent protein for cellular imaging. Nat Methods 14:53–56.

81. Datsenko KA, Wanner BL. 2000. One-step inactivation of chromosomal genes in Escherichia coli K-12 using PCR products. Proc Natl Acad Sci U S A 97:6640–6645.

82. Kanda T, Abiko G, Kanesaki Y, Yoshikawa H, Iwai N, Wachi M. 2020. RNase E-dependent degradation of tnaA mRNA encoding tryptophanase is prerequisite for the induction of acid resistance in Escherichia coli. Sci Rep 10.

83. Blank K, Hensel M, Gerlach RG. 2011. Rapid and highly efficient method for scarless mutagenesis within the salmonella enterica chromosome. PLoS One 6.

84. Uzzau S, Figueroa-Bossi N, Rubino S, Bossi L. 2001. Epitope tagging of chromosomal genes in Salmonella. Proc Natl Acad Sci U S A 98:15264–15269.

85. Papenfort K, Sun Y, Miyakoshi M, Vanderpool CK, Vogel J. 2013. Small RNA-mediated activation of sugar phosphatase mRNA regulates glucose homeostasis. Cell 153:426–437.

86. Livak KJ, Schmittgen TD. 2001. Analysis of relative gene expression data using real-time quantitative PCR and the 2-ΔΔCT method. Methods 25:402–408.

87. Chao Y, Papenfort K, Reinhardt R, Sharma CM, Vogel J. 2012. An atlas of Hfq-bound transcripts reveals 3′ UTRs as a genomic reservoir of regulatory small RNAs. EMBO Journal 31:4005–4019.

88. Melamed S, Faigenbaum-Romm R, Peer A, Reiss N, Shechter O, Bar A, Altuvia Y, Argaman L, Margalit H. 2018. Mapping the small RNA interactome in bacteria using RIL-seq. Nat Protoc 13:1–33.

89. Katoh K, Rozewicki J, Yamada KD. 2018. MAFFT online service: Multiple sequence alignment, interactive sequence choice and visualization. Brief Bioinform 20:1160–1166.

90. Corpet F. 1988. Multiple sequence alignment with hierarchical clustering. Nucleic Acids Res 16:10881–10890.

