## Supplementary material for "3′UTR-derived small RNA couples acid resistance to metabolic reprogramming of *Salmonella* within macrophages": Table_S3_S4_S5

**Table S3. Bacterial and virus strains used in this study.**

| Bacterial and virus strains | Source or reference |
|---|---|
| *Salmonella Typhimurium* SL1344 | Lab stock |
| *Salmonella* Δ*adiA*::$Km^R$ | This work |
| *Salmonella* Δ*adiZ*::$Km^R$ | This work |
| *Salmonella* Δ*adiZ*::FRT | This work |
| *Salmonella* Δ*adiAZ*::$Km^R$ | This work |
| *Salmonella* Δ*adiY*::$Km^R$ | This work |
| *Salmonella* Δ*adiZ*::$Cm^R$ I-SceI | This work |
| *Salmonella pykF*-3xFLAG-$Km^R$ | This work |
| *Salmonella* Δ*adiZ*::FRT *pykF*-3xFLAG-$Km^R$ | This work |
| *Salmonella adiZ*-G10C *pykF*-3xFLAG-$Km^R$ | This work |
| *Salmonella adiZ*-G32C *pykF*-3xFLAG-$Km^R$ | This work |
| *Salmonella adiZ*-C46G *pykF*-3xFLAG-$Km^R$ | This work |
| JVS-4141 | (1) |
| *Salmonella ptsG*-3xFLAG-$Km^R$ | This work |
| *Salmonella* Δ*adiZ*::FRT *ptsG*-3xFLAG-$Km^R$ | This work |
| *Salmonella adiZ*-G10C *ptsG*-3xFLAG-$Km^R$ | This work |
| *Salmonella adiZ*-G32C *ptsG*-3xFLAG-$Km^R$ | This work |
| *Salmonella adiZ*-C46G *ptsG*-3xFLAG-$Km^R$ | This work |
| *Salmonella dmsA*-3xFLAG-$Km^R$ | This work |
| *Salmonella* Δ*adiZ*::FRT *dmsA*-3xFLAG-$Km^R$ | This work |
| *Salmonella adiZ*-G10C *dmsA*-3xFLAG-$Km^R$ | This work |
| *Salmonella adiZ*-G32C *dmsA*-3xFLAG-$Km^R$ | This work |
| *Salmonella adiZ*-C46G *dmsA*-3xFLAG-$Km^R$ | This work |
| *Salmonella* $hisG^+$ | This work |
| *Salmonella* $hisG^+$ *putAP*::$P_{Llac\text{-}O}$-$tnaC_{eco}$5'UTR-sfGFP-$Cm^R$ | This work |
| *Salmonella* $hisG^+$ *putAP*::$P_{Llac\text{-}O}$-$tnaC_{eco}$5'UTR-mSc-$Cm^R$ | This work |
| *Salmonella* Δ*adiZ*::FRT $hisG^+$ *putAP*::$P_{Llac\text{-}O}$-$tnaC_{eco}$5'UTR-sfGFP-$Cm^R$ | This work |
| *Salmonella* Δ*adiZ*::FRT $hisG^+$ *putAP*::$P_{Llac\text{-}O}$-$tnaC_{eco}$5'UTR-mSc-$Cm^R$ | This work |
| *Salmonella adiZ*-G10C $hisG^+$ *putAP*::$P_{Llac\text{-}O}$-$tnaC_{eco}$5'UTR-mSc-$Cm^R$ | This work |
| *Salmonella adiZ*-G32C $hisG^+$ *putAP*::$P_{Llac\text{-}O}$-$tnaC_{eco}$5'UTR-mSc-$Cm^R$ | This work |
| *Salmonella adiZ*-C46G $hisG^+$ *putAP*::$P_{Llac\text{-}O}$-$tnaC_{eco}$5'UTR-mSc-$Cm^R$ | This work |
| *Salmon*ella *hfq*::3xFLAG-$Km^R$ | (2) |
| *E. coli* MG1655 | Lab stock |

| | |
|---|---|
| *E. coli* Δ*adiZ*::$Km^R$ | This work |
| *E. coli* Δ*adiZ*::FRT | This work |
| *E. coli* Δ*hfq*::$Km^R$ | This work |
| *E. coli* Δ*adiZ*::FRT Δ*hfq*::$Km^R$ | This work |
| TK40 | (3) |
| *E. coli* Δ*adiZ*::FRT *rne-1* | This work |
| *E. coli* DH5α | NIPPON GENE |
| Bacteriophage P1 vir | (4) |
| Bacteriophage P22 HT105/1 *int-201* | (5) |

### Table S4. Plasmids used in this study.

| Name | Relevant fragment | Comment | Origin / marker | Reference |
|---|---|---|---|---|
| pKP8-35 | Control plasmid | pBAD control plasmid, expresses ~50 nt nonsense RNA derived from *rrnB* terminator | pBR322 / $Amp^R$ | (6) |
| pBAD-AdiAZ | $P_{BAD}$-*adiAZ* | *Salmonella adiA-adiZ* mid-copy expression plasmid, *adiA-adiZ* is controlled by the L-arabinose-inducible $P_{BAD}$ promoter | pBR322 / $Amp^R$ | This work |
| pBAD-AdiZ | $P_{BAD}$-*adiZ* | *Salmonella adiZ* mid-copy expression plasmid, *adiZ* is controlled by the L-arabinose-inducible $P_{BAD}$ promoter | pBR322 / $Amp^R$ | This work |
| pYC582 | $P_{BAD}$-*t4rnl1* | T4 RNA ligase 1 expression plasmid, *t4rnl1* is controlled by the L-arabinose-inducible $P_{BAD}$ promoter | pBR322 / $HyG^R$ | (2) |
| pJV300 | Control plasmid | $pP_L$ control plasmid, expresses ~50 nt nonsense RNA derived from *rrnB* terminator | ColE1 / $Amp^R$ | (7) |
| $pP_L$-AdiZ | $P_{Llac\text{-}O}$-*adiZ* | *Salmonella adiZ* mid-copy expression plasmid, *adiZ* is controlled by the constitutive $P_{LlacO}$ promoter | ColE1 / $Amp^R$ | This work |
| $pP_L$-AdiZ G10C | $P_{Llac\text{-}O}$-*adiZ* G10C | *Salmonella adiZ* mutant in position 10 (G->C) | ColE1 / $Amp^R$ | This work |
| $pP_L$-AdiZ G32C | $P_{Llac\text{-}O}$-*adiZ* G32C | *Salmonella adiZ* mutant in position 32 (G->C) | ColE1 / $Amp^R$ | This work |
| $pP_L$-AdiZ C46G | $P_{Llac\text{-}O}$-*adiZ* C46G | *Salmonella adiZ* mutant in position 46 (C->G) | ColE1 / $Amp^R$ | This work |
| pXG-10sf | $P_{LtetO}$- *lacZ*::*gfp* | Plasmid for construction of translational sfGFP fusion | pSC101* / $Cm^R$ | (8) |
| pXG-30sf | $P_{LtetO}$-FLAG::*glmU-glmS*::*gfp* | Plasmid for construction of translational sfGFP fusions of dicistronic targets | pSC101* / $Cm^R$ | (8) |
| pXG-10sf-*pykF* | $P_{LtetO}$-*pykF*::*gfp* | *Salmonella pykF* translational GFP fusion plasmid | pSC101*/ $Cm^R$ | This work |
| pXG-10sf-$pykF_{C\text{-}22G}$ | $P_{LtetO}$-$pykF_{C\text{-}22G}$::*gfp* | *Salmonella pykF* mutant in position -22 relative to the start codon (C->G) | pSC101*/ $Cm^R$ | This work |
| pXG-10sf-*ptsG* | $P_{LtetO}$-*ptsG*::*gfp* | *Salmonella ptsG* translational GFP fusion plasmid | pSC101*/ $Cm^R$ | This work |
| pXG-10sf-$ptsG_{C\text{-}19G}$ | $P_{LtetO}$-$ptsG_{C\text{-}19G}$::*gfp* | *Salmonella ptsG* mutant in position -19 relative to the start codon (C->G) | pSC101*/ $Cm^R$ | This work |
| pXG-10sf-*dmsA* | $P_{LtetO}$-*dmsA*::*gfp* | *Salmonella dmsA* translational GFP fusion plasmid | pSC101*/ $Cm^R$ | This work |
| pXG-10sf-$dmsA_{G\text{-}22C}$ | $P_{LtetO}$-$dmsA_{G\text{-}22C}$::*gfp* | *Salmonella dmsA* mutant in position -22 relative to the start codon (G->C) | pSC101*/ $Cm^R$ | This work |
| pXG-10sf-*cadB* | $P_{LtetO}$-*cadB*::*gfp* | *Salmonella cadB* translational GFP fusion plasmid | pSC101*/ $Cm^R$ | This work |
| pXG-10sf-*tktA* | $P_{LtetO}$-*tktA*::*gfp* | *Salmonella tktA* translational GFP fusion plasmid | pSC101*/ $Cm^R$ | This work |
| pXG-10sf-*yqhD* | $P_{LtetO}$-*yqhD*::*gfp* | *Salmonella yqhD* translational GFP fusion plasmid | pSC101*/ $Cm^R$ | This work |
| pXG-10sf-*dkgA* | $P_{LtetO}$-*dkgA*::*gfp* | *Salmonella dkgA* translational GFP fusion plasmid | pSC101*/ $Cm^R$ | This work |
| pXG-10sf-RS16625 | $P_{LtetO}$-RS16625::*gfp* | *Salmonella* RS16625 translational GFP fusion plasmid | pSC101*/ $Cm^R$ | This work |
| pXG-10sf-*caiF* | $P_{LtetO}$-*caiF*::*gfp* | *Salmonella caiF* translational GFP fusion plasmid | pSC101*/ $Cm^R$ | This work |
| pXG-10sf-*adiY* | $P_{LtetO}$-*adiY*::*gfp* | *Salmonella adiY* translational GFP fusion plasmid | pSC101*/ $Cm^R$ | This work |
| pXG-10sf-*fimA* | $P_{LtetO}$-*fimA*::*gfp* | *Salmonella fimA* translational GFP fusion plasmid | pSC101*/ $Cm^R$ | This work |
| pXG-10sf-*fimI* | $P_{LtetO}$-*fimI*::*gfp* | *Salmonella fimI* translational GFP fusion plasmid | pSC101*/ $Cm^R$ | This work |

| | | | | |
|---|---|---|---|---|
| pXG-10sf-*cadC* | $P_{LtetO}$-*cadC*::*gfp* | *Salmonella cadC* translational GFP fusion plasmid | pSC101*/ $Cm^R$ | This work |
| pXG-10sf-*lrhA* | $P_{LtetO}$-*lrhA*::*gfp* | *Salmonella lrhA* translational GFP fusion plasmid | pSC101*/ $Cm^R$ | This work |
| pXG-10sf-*hdeB* | $P_{LtetO}$-*hdeB*::*gfp* | *Salmonella hdeB* translational GFP fusion plasmid | pSC101*/ $Cm^R$ | This work |
| pXG-10sf-*araJ* | $P_{LtetO}$-*araJ*::*gfp* | *Salmonella araJ* translational GFP fusion plasmid | pSC101*/ $Cm^R$ | This work |
| pXG-10sf-*yfbV* | $P_{LtetO}$-*yfbV*::*gfp* | *Salmonella yfbV* translational GFP fusion plasmid | pSC101*/ $Cm^R$ | (9) |
| pXG-10sf-*trxA* | $P_{LtetO}$-*trxA*::*gfp* | *Salmonella trxA* translational GFP fusion plasmid | pSC101*/ $Cm^R$ | This work |
| pXG-30sf-*cadB-cadA* | $P_{LtetO}$-FLAG::*cadB-cadA*::*gfp* | *Salmonella cadB-cadA* translational GFP fusion plasmid | pSC101*/ $Cm^R$ | this study |
| pXG-30sf-*atpG-atpD* | $P_{LtetO}$-FLAG::*atpG-atpD*::*gfp* | *Salmonella atpG-atpD* translational GFP fusion plasmid | pSC101*/ $Cm^R$ | this study |
| pXG-30sf-*tolB-pal* | $P_{LtetO}$-FLAG::*tolB-pal*::*gfp* | *Salmonella tolB-pal* translational GFP fusion plasmid | pSC101*/ $Cm^R$ | this study |
| pXG-10sf-$tnaC_{eco}$ | $P_{LtetO}$-$tnaC_{eco}$::*gfp* | *E. coli tnaC* translational GFP fusion plasmid | pSC101*/ $Cm^R$ | (10) |
| pXG-10-$tnaC_{eco}$5'UTR-sfGFP | $P_{LtetO}$-$tnaC_{eco}$5'UTR::*gfp* | *E. coli tnaC* 5'UTR-GFP fusion plasmid | pSC101*/ $Cm^R$ | this study |
| pXG-10-$tnaC_{eco}$5'UTR-mSc | $P_{LtetO}$-$tnaC_{eco}$5'UTR::*mSc* | *E. coli tnaC* 5'UTR-mSc fusion plasmid | pSC101*/ $Cm^R$ | this study |
| pCWU6-mCherry | | Template of mCherry | pBR322 / $Amp^R$ | (7) |
| pYC963 | $P_{Llac\text{-}O}$-*adiZ* -mCherry | mCherry mid-copy expression plasmid, mCherry is controlled by the constitutive $P_{LlacO}$ promoter | pBR322 / $Amp^R$ | This work |
| pYC964 | $P_{adiA}$-mCherry | mCherry mid-copy expression plasmid, mCherry is controlled by the $P_{adiA}$ promoter | pBR322 / $Amp^R$ | This work |
| pKD13 | | Template of $Km^R$ cassette | oriRγ / $Amp^R$ | (11) |
| pKD46 | | Temperature-sensitive λ Red expression plasmid | oriR101 / $Amp^R$ | (11) |
| pCP20 | | Temperature-sensitive FLP expression plasmid | oriR101 / $Amp^R$ | (11) |
| pWRG99 | | Temperature-sensitive λ Red and I-SceI expression plasmid | oriR101 / $Amp^R$ | (12) |
| pWRG100 | | Template of $Cm^R$ I-SceI cassette | oriRγ / $Amp^R$ | (12) |

### Table S5. DNA oligonucleotides used in this study.

| Name | Sequence (5' -> 3' direction) | Purpose |
|---|---|---|
| **AdiZ cloning** | | |
| MMO-1496 | ACATTCATACGGCGGGTCG | Sense oligo for *adiA-adiZ* cloning into the pBAD or $pP_L$ plasmid |
| MMO-0543 | ATCACAAAGAGTCTTTTTTTTGAGG | Sense oligo for AdiZ cloning into the pBAD or $pP_L$ plasmid |
| MMO-0544 | GTTTTTTCTAGAGTTGTACGCGAGTGGAAAC | Antisense oligo for *adiA-adiZ* or AdiZ cloning into the pBAD or $pP_L$ plasmid |
| **AdiZ mutagenesis on $pP_L$-AdiZ** | | |
| MMO-1933 | TTTTTTCAGGATGGCATGTATTCC | Sense oligo for AdiZ G10C mutation |
| MMO-1934 | CATCCTGAAAAAAAAGACTCTTTGTG | Antisense oligo for AdiZ G10C mutation |
| MMO-2095 | CCCTGACCTATTGCTGCAAAACAATAATAC | Sense oligo for AdiZ G32C mutation |
| MMO-2096 | GCAATAGTCAGGGAATACATGCCATC | Antisense oligo for AdiZ G32C mutation |
| MMO-1807 | GCAAAAGAATAATACTGTCATACCTTTTTG | Sense oligo for AdiZ C46G mutation |
| MMO-1808 | ATTATTCTTTTGCAGCAATACTCAGG | Antisense oligo for AdiZ C46G mutation |
| **AdiZ target candidate cloning** | | |
| MMO-1817 | GTTTTTATGCATAATTGTCTCTTGAATGGTTTCAGC | Sense oligo for *pykF* GFP fusion cloning |
| MMO-1818 | GTTTTTGCTAGCGTCCAGCATTTTGCTTAACATCTC | Antisense oligo for *pykF* GFP fusion cloning |
| MMO-2050 | GTTTTTATGCATAATAAATAAAAGGGCACTTAGATGTCC | Sense oligo for *ptsG* GFP fusion cloning |
| MMO-2051 | GTTTTTGCTAGCTGCATTCTTAAACATAATTGAGAGTG | Antisense oligo for *ptsG* GFP fusion cloning |
| MMO-1799 | GTTTTTATGCATGAATGAAGGTCAGTTTTTAACAC | Sense oligo for *dmsA* GFP fusion cloning |
| MMO-1800 | GTTTTTGCTAGCTATCGCGGTGGTTTTGAC | Antisense oligo for *dmsA* GFP fusion cloning |
| MMO-1819 | GTTTTTATGCATAGTTGAACGGGCACGCAGC | Sense oligo for *cadB* GFP fusion cloning |
| MMO-1820 | GTTTTTGCTAGCTAGGCCAATCGTTGTCAAGCGG | Antisense oligo for *cadB* GFP fusion cloning |
| MMO-1821 | GTTTTTATGCATAAATTTTCCGGCGTAGCCCAAAACG | Sense oligo for *tktA* GFP fusion cloning |
| MMO-1822 | GTTTTTGCTAGCATTGGACAGCACGAAGCGGTC | Antisense oligo for *tktA* GFP fusion cloning |
| MMO-1795 | GTTTTTATGCATCATCAGGCAGACAGCCGCC | Sense oligo for *yqhD* GFP fusion cloning |
| MMO-1796 | GTTTTTGCTAGCCTGCGGGATTTGATCGCGAAG | Antisense oligo for *yqhD* GFP fusion cloning |
| MMO-1793 | GTTTTTATGCATACTTAGTCTCGCCGGACCC | Sense oligo for *dkgA* GFP fusion cloning |
| MMO-1794 | GTTTTTGCTAGCGGTATCAATCGATCGATAGCCCAC | Antisense oligo for *dkgA* GFP fusion cloning |
| MMO-1825 | GTTTTTATGCATGTATGAGGATATGCAATCCCAGGG | Sense oligo for RS16625 GFP fusion cloning |
| MMO-1826 | GTTTTTGCTAGCAGACAGAGCGGAACTCACCAC | Antisense oligo for RS16625 GFP fusion cloning |
| MMO-1823 | GTTTTTATGCATATATCCACAATTTTAATATGGCTTAG | Sense oligo for *caiF* GFP fusion cloning |
| MMO-1824 | GTTTTTGCTAGCGCGACGCATACTCTTGGAG | Antisense oligo for *caiF* GFP fusion cloning |

| | | |
|---|---|---|
| MMO-1801 | GTTTTTATGCATGTTTCAATTCATCACACAATTGACTC | Sense oligo for *adiY* GFP fusion cloning |
| MMO-1802 | GTTTTTGCTAGCTGGCTCCTTACCATTTACTCG | Antisense oligo for *adiY* GFP fusion cloning |
| MMO-1780 | GTTTTTATGCATGTGACGAAATGTCATATTCGCAAAG | Sense oligo for *fimA* GFP fusion cloning |
| MMO-1782 | GTTTTTGCTAGCCTGCGCAGTCGTATTACCAATC | Antisense oligo for *fimA* GFP fusion cloning |
| MMO-1797 | GTTTTTATGCATGGCTATCGGGGATAAATAGAGAAC | Sense oligo for *fimI* GFP fusion cloning |
| MMO-1798 | GTTTTTGCTAGCGTGAACACGCCCGCTCTC | Antisense oligo for *fimI* GFP fusion cloning |
| MMO-1918 | GTTTTTATGCATTTTGACTTAGCTCGTTAGGGC | Sense oligo for *cadC* GFP fusion cloning |
| MMO-1919 | GTTTTTGCTAGCTGGTTCAAGAGTAATCTGGCG | Antisense oligo for *cadC* GFP fusion cloning |
| MMO-2158 | GTTTTTATGCATCGAAACGGGGGCGTAGTC | Sense oligo for *lrhA* GFP fusion cloning |
| MMO-2159 | GTTTTTGCTAGCCAGATCGAGATCGAGGTTAATTATCGG | Antisense oligo for *lrhA* GFP fusion cloning |
| MMO-2048 | GTTTTTATGCATGTTCTACATTAACGGGTTAACGCG | Sense oligo for *hdeB* GFP fusion cloning |
| MMO-2049 | GTTTTTGCTAGCTTTAGTAGTATCTGTTGCCGCATTC | Antisense oligo for *hdeB* GFP fusion cloning |
| MMO-2046 | GTTTTTATGCATGAATACACAAAGACGGATAAGGAATGG | Sense oligo for *araJ* GFP fusion cloning |
| MMO-2047 | GTTTTTGCTAGCAATAACTTTTTTCATACCACCTGCC | Antisense oligo for *araJ* GFP fusion cloning |
| MMO-2052 | GTTTTTATGCATAACCTTTGTTGGCGAAGTTAAC | Sense oligo for *trxA* GFP fusion cloning |
| MMO-2053 | GTTTTTGCTAGCAATTTTATCGCTCATATATAACTCC | Antisense oligo for *trxA* GFP fusion cloning |
| MMO-1920 | GTTTTTATGCATGTCAGTCTGATCATCCTGATGTTC | Sense oligo for *cadB-cadA* GFP fusion cloning |
| MMO-1921 | GTTTTTGCTAGCATCCCAGTCGAAAATGACGC | Antisense oligo for *cadB-cadA* GFP fusion cloning |
| MMO-1587 | GTTTTTATGCATCTGATTAAAGAGCTGCAGTTGG | Sense oligo for *atpG-atpD* GFP fusion cloning |
| MMO-1588 | GTTTTTGCTAGCCACCAGCTTCTCATTACCATTC | Antisense oligo for *atpG-atpD* GFP fusion cloning |
| MMO-1627 | GTTTTTATGCATGGCGTACAGGTTCTGTCGTC | Sense oligo for *tolB-pal* GFP fusion cloning |
| MMO-1628 | GTTTTTGCTAGCGCCGTTAGCGTCCATACCAG | Antisense oligo for *tolB-pal* GFP fusion cloning |
| **AdiZ target mutagenesis** | | |
| MMO-1935 | CCTCCTGAACTTAAAGACTAAGACTG | Sense oligo for *pykF* C-22G mutation |
| MMO-1936 | TAAGTTCAGGAGGATATGGAAATCTG | Antisense oligo for *pykF* C-22G mutation |
| MMO-2093 | CAAATAGTCAGGAGCACTCTCAATTATG | Sense oligo for *ptsG* C-19G mutation |
| MMO-2094 | TCCTGACTATTTGTGCTTTTCTACG | Antisense oligo for *ptsG* C-19G mutation |
| MMO-1815 | TTTATTCTGAGCAGCAGAGTGAG | Sense oligo for *dmsA* G-22C mutation |
| MMO-1816 | TGCTCAGAATAAAAAAACAATAAAACGATG | Antisense oligo for *dmsA* G-22C mutation |
| **λ Red recombination** | | |
| MMO-2215 | GCAAGGCGTAAATTGCACGGCCTCCACAACCGGGTAAAAGCTG TCAAACATGAGAATTAATTC | Sense oligo to construct *Salmonella* Δ*adiA*::Km$^R$ or Δ*adiAZ*::Km$^R$ with pKD13 |
| MMO-2216 | CAGGGAATACATGCCATCCTCAAAAAAAAGACTCTTTGTGAGT GTAGGCTGGAGCTGCTTC | Antisense oligo to construct *Salmonella* Δ*adiA*::Km$^R$ with pKD13 |

| | | |
|---|---|---|
| MMO-1486 | GCACGGAAATTATTGATGGCGTATATCACGTTATGTGCGTGAAAGCCTAACTGTCAAACATGAGAATTAATTC | Sense oligo to construct *Salmonella* Δ*adiZ*::Km$^R$ with pKD13 |
| MMO-1487 | GTGGAAACTTGTCATTTTTGTATAAAAAAAGCGCGGGATGGGGCCCGCGCGTGTAGGCTGGAGCTGCTTC | Antisense oligo to construct *Salmonella* Δ*adiAZ*::Km$^R$ or Δ*adiZ*::Km$^R$ with pKD13 |
| MMO-2162 | GTTTGGCGAGATCCGGTCTTTTATGTTCATATTCCAGGAGTTCGGGTATGCTGTCAAACATGAGAATTAATTC | Sense oligo to construct *Salmonella* Δ*adiY*::Km$^R$ with pKD13 |
| MMO-2163 | ATCGTGTCTTATCGTTGAGTGTAGAAAAATTAGGCAGCCGATCGTTCCCGGTGTAGGCTGGAGCTGCTTC | Antisense oligo to construct *Salmonella* Δ*adiY*::Km$^R$ with pKD13 |
| MMO-2217 | CACTTACATTTGTTGCGAACCTTTGGGAGTACAGACAATGCTGTCAAACATGAGAATTAATTC | Sense oligo to construct *Salmonella* Δ*ompR*::Km$^R$ with pKD13 |
| MMO-2218 | GGTGAGAAGCGCATTCGCCTCATGCTTTAGAACCGTCCGGGTGTAGGCTGGAGCTGCTTC | Antisense oligo to construct *Salmonella* Δ*ompR*::Km$^R$ with pKD13 |
| MMO-1484 | GGACTGAAATTATTGACGGTATTTACCACGTTATGTGCGTGAAAGCGTAACTGTCAAACATGAGAATTAATTC | Sense oligo to construct *E. coli* Δ*adiZ*::Km$^R$ with pKD13 |
| MMO-1485 | TTTTTACAGGATAAATAACAGGCAAAAAAAAGCGCGGGGTGGGGCCGCGCGTGTAGGCTGGAGCTGCTTC | Antisense oligo to construct *E. coli* Δ*adiZ*::Km$^R$ with pKD13 |
| hfqDeletion-F1 | TCGAAAGGTTCAAAGTACAAATAAGCATATAAGGAAAAGAGAGAATGCTGTCAAACATGAGAATTAATTC | Sense oligo to construct *E. coli* Δ*hfq*::Km$^R$ with pKD13 |
| hfqDeletion-R1 | CTCCCCGTGTAAAAAAACAGCCCGAAACCTTATTCGGTTTCTTCGCTGTCGTGTAGGCTGGAGCTGCTTC | Antisense oligo to construct *E. coli* Δ*hfq*::Km$^R$ with pKD13 |
| MMO-2126 | GCACGGAAATTATTGATGGCGTATATCACGTTATGTGCGTGAAAGCCTAAGTGTAGGCTGGAGCTGCTTC | Sense oligo to construct *Salmonella* Δ*adiZ*::Cm$^R$ I-SceI with pWRG100 |
| MMO-2127 | GTGGAAACTTGTCATTTTTGTATAAAAAAAGCGCGGGATGGGGCCCGCGCCTAGACTATATTACCCTGTT | Antisense oligo to construct *Salmonella* Δ*adiZ*::Cm$^R$ I-SceI with pWRG100 |
| MMO-2129 | TTATGTGCGTGAAAGCCTAATCACAAAGAGTCTTTTTTTTCAGG | Sense oligo for AdiZ-G10C cloning with pPL-AdiZ G10C |
| MMO-2128 | TTATGTGCGTGAAAGCCTAATCACAAAGAGTCTTTTTTTTGAGG | Sense oligo for AdiZ-G32C or C46G cloning with pPL-AdiZ G32C or C46G |
| MMO-2130 | GTTGTACGCGAGTGGAAAC | Antisense oligo for AdiZ cloning with pPL-AdiZ derivatives |
| MMO-2146 | TCCAAGCGGCACCACTAACACCGCCTCTGTTCACGTACTGGACTACAAAGACCATGACGG | Sense oligo to construct *Salmonella pykF*-3xFLAG-Km$^R$ with pSUB11 |
| MMO-2147 | AAGGGCGCTTTTTTAAACAAATTAATTCACACAACAATTACATATGAATATCCTCCTTAGTTCC | Antisense oligo to construct *Salmonella pykF*-3xFLAG-Km$^R$ with pSUB11 |
| MMO-2148 | GAACCCGTCACATACGAACCTCGTTCAGGTTGAAAAGGCGGACTACAAAGACCATGACGG | Sense oligo to construct *Salmonella dmsA*-3xFLAG-Km$^R$ with pSUB11 |
| MMO-2149 | ATCAATAAAAAATCCATACTGGGTTGTCATCGGTTACTCCCATATGAATATCCTCCTTAGTTCC | Antisense oligo to construct *Salmonella dmsA*-3xFLAG-Km$^R$ with pSUB11 |
| ***hisG* reverse mutation** | | |
| MMO-1943 | ACACGCGTTCAATTTAAACACC | Sense oligo for *hisG* cloning |
| MMO-1944 | CGTGGTCGATCTCGGTATTATCG | Sense oligo for *hisG* mutation (C206T) |
| MMO-1945 | TCAGACAGGCTTTAGAGCGG | Antisense oligo for *hisG* cloning |
| MMO-1946 | CGATAATACCGAGATCGACCACG | Antisense oligo for *hisG* mutation (C206T) |
| **AdiA transcriptional reporter** | | |
| YCO-3283 | AACACTTTCGTCGGGGATTCG | Sense oligo for amplifying pZE12 backbone |
| YCO-3284 | ACGATAAGGCAAATGACGATTAAGCC | Antisense oligo for amplifying pZE12 backbone |
| YCO-3285 | CATTTGCCTTATCGTGCTAGCGTGAGCAAGGGCGAGGAG | Sense oligo for amplifying mCherry |
| YCO-3286 | CCCGACGAAAGTGTTCTACTTGTACAGCTCGTCCATGCC | Antisense oligo for amplifying mCherry |
| YCO-3367 | CTTGCTCACGCTAGCGGTGTCCTGATGCAGAAACTCAC | Sense oligo for amplifying pZE12-mCherry backbone |
| YCO-3368 | AATAGGGGTTCCGCGGCCGGATACCACTATCAGCCAA | Antisense oligo for amplifying pZE12-mCherry backbone |
| YCO-3369 | CTTGCTCACGCTAGCGGTGTCCTGATGCAGAAACTCAC | Sense oligo for amplifying *adiA* promoter |

| | | |
|---|---|---|
| YCO-3370 | AATAGGGGTTCCGCGGCCGGATACCACTATCAGCCAA | Antisense oligo for amplifying *adiA* promoter |
| **Fluorescence marker insertion into *Salmonella* chromosome** | | |
| MMO-1937 | CATAACTTGCTCCTTAGCCGTTATC | Sense oligo for sfGFP on pXG-10/30sf |
| MMO-1987 | CATAATGCACTTATCCTCGCAAGAC | Antisense oligo for *tnaC*$_{eco}$ on pXG-10sf-*tnaC*$_{eco}$ |
| MMO-1982 | AATGACTCTAGAGGCATCAAATAAAACGAAAGG | Sense oligo for just downstream of sfGFP on pXG-10/30sf |
| MMO-2000 | GTTTTTGCTAGCCATAATGCACTTATCCTCGCAAGAC | Antisense oligo for 5'UTR-start codon of *tnaC*$_{eco}$ on pXG-10sf-*tnaC*$_{eco}$ |
| MMO-1998 | GTTTTTGCTAGCGTATCTAAAGGTGAAGCTGTTATTAAAGAG | Sense oligo for mSc |
| MMO-1999 | GTTTTTTCTAGATTACTTATAAAGTTCATCCATCCCTCC | Antisense oligo for mSc |
| MMO-1988 | CTCCTGTTATTCGCTCTATCTGCTAACCAATAGTTAGCGGAAAATATCCAATTCTCACCAATAAAAAACGCCCG | Sense oligo to construct *Salmonella putAP*::P$_{Llac\text{-}O}$-*tnaC*$_{eco}$5'UTR-sfGFP/mSc-Cm$^R$ with pXG-10-*tnaC*$_{eco}$5'UTR-sfGFP/mSc |
| MMO-1989 | GCTTGCAACGCAGGTCACATTTAACGCGGTTGCACAAAGTTGCAACATGGATTTGTCCTACTCAGGAGAGCG | Antisense oligo to construct *Salmonella putAP*::P$_{Llac\text{-}O}$-*tnaC*$_{eco}$5'UTR-sfGFP/mSc-Cm$^R$ with pXG-10-*tnaC*$_{eco}$5'UTR-sfGFP/mSc |
| **In vitro transcription** | | |
| AWO-1787 | GTTTTTTTTAATACGACTCACTATAGGCTTTTTTTTGAGGATGGCATGTATTC | Sense oligo to amplify *adiZ* template with T7 promoter for IVT |
| AWO-1788 | AAAAAAAGCGCGGGATGG | Antisense oligo to amplify *adiZ* template |
| AWO-1801 | GTTTTTTTTAATACGACTCACTATAGGGAGGCATCATCACTTTAGCAACAC | Sense oligo to amplify *pykF* template with T7 promoter for IVT |
| AWO-1802 | CATGACAGTCTTAGTCTTTAAGTTGAGG | Antisense oligo to amplify *pykF* template |
| AWO-1803 | GTTTTTTTTAATACGACTCACTATAGGAATAAATAAAAGGGCACTTAGATGTCC | Sense oligo to amplify *ptsG* template with T7 promoter for IVT |
| AWO-1804 | TGCATTCTTAAACATAATTGAGAGTG | Antisense oligo to amplify *ptsG* template |
| AWO-1805 | GTTTTTTTTAATACGACTCACTATAGGGAATGAAGGTCAGTTTTTAACAC | Sense oligo to amplify *dmsA* template |
| AWO-1806 | CATAAGGGCTCACTCTGC | Antisense oligo to amplify *dmsA* template |
| **Northern blot** | | |
| MMO-1468 | ATGAAAACGTTGCCATACTC | *E. coli* AdiZ oligo probe |
| MMO-1469 | AGGGAATACATGCCATCCTC | *Salmonella* AdiZ oligo probe, complementary to 10-29 nt |
| MMO-2143 | TCAGGGAATACATGCCATCC | *Salmonella* AdiZ oligo probe, complementary to 12-31 nt |
| MMO-1056 | ACTACCATCGGCGCTACGGC | 5S rRNA oligo probe |
| **qRT-PCR** | | |
| MMO-1759 | TCCTTTCTATCCAGCTTGACTCG | Sense oligo for quantification of *rpoB* mRNA levels |
| MMO-1760 | TCGGAATTACCGCTGTAGCTC | Antisense oligo for quantification of *rpoB* mRNA levels |
| MMO-1941 | TTCTGCATCAGGACACCTGG | Sense oligo for quantification of *adiA* mRNA levels |
| MMO-1942 | TCGCGGACAAGATGGCATAG | Antisense oligo for quantification of *adiA* mRNA levels |
| MMO-2166 | GAGCCAATCAGCTTAAAAGCG | Sense oligo for quantification of *adiY* mRNA levels |
| MMO-2167 | GCGACTCACTTGTATTACTAGCAC | Antisense oligo for quantification of *adiY* mRNA levels |

XbaI, NsiI, and NheI sites used for cloning are highlighted in yellow, cyan, and green, respectively.

Point mutations introduced by site-directed mutagenesis are highlighted in magenta.

**References**

1. Papenfort K, Sun Y, Miyakoshi M, Vanderpool CK, Vogel J. 2013. Small RNA-mediated activation of sugar phosphatase mRNA regulates glucose homeostasis. Cell 153:426–437.

2. Liu F, Chen Z, Zhang S, Wu K, Bei C, Wang C, Chao Y. 2023. In vivo RNA interactome profiling reveals 3'UTR-processed small RNA targeting a central regulatory hub. Nat Commun 14.

3. Kanda T, Abiko G, Kanesaki Y, Yoshikawa H, Iwai N, Wachi M. 2020. RNase E-dependent degradation of tnaA mRNA encoding tryptophanase is prerequisite for the induction of acid resistance in Escherichia coli. Sci Rep 10.

4. Ikeda H, Tomizawa J ichi. 1965. Transducing fragments in generalized transduction by phage P1: I. Molecular origin of the fragments. J Mol Biol 14:85–109.

5. Schmieger H. 1972. Phage P22-mutants with increased or decreased transduction abilities. MGG Molecular & General Genetics 119:75–88.

6. Papenfort K, Pfeiffer V, Mika F, Lucchini S, Hinton JCD, Vogel J. 2006. SigmaE-dependent small RNAs of Salmonella respond to membrane stress by accelerating global omp mRNA decay. Mol Microbiol 62:1674–1688.

7. Sittka A, Pfeiffer V, Tedin K, Vogel J. 2007. The RNA chaperone Hfq is essential for the virulence of Salmonella typhimurium. Mol Microbiol 63:193–217.

8. Corcoran CP, Podkaminski D, Papenfort K, Urban JH, Hinton JCD, Vogel J. 2012. Superfolder GFP reporters validate diverse new mRNA targets of the classic porin regulator, MicF RNA. Mol Microbiol 84:428–445.

9. Miyakoshi M, Matera G, Maki K, Sone Y, Vogel J. 2019. Functional expansion of a TCA cycle operon mRNA by a 3 end-derived small RNA. Nucleic Acids Res 47:2075–2088.

10. Kanda T, Sekijima T, Miyakoshi M. 2025. Post-transcriptional regulation of aromatic

amino acid metabolism by GcvB small RNA in Escherichia coli. Microbiol Spectr 13.

11. Datsenko KA, Wanner BL. 2000. One-step inactivation of chromosomal genes in Escherichia coli K-12 using PCR products. Proc Natl Acad Sci U S A 97:6640–6645.

12. Blank K, Hensel M, Gerlach RG. 2011. Rapid and highly efficient method for scarless mutagenesis within the salmonella enterica chromosome. PLoS One 6.
